# A harmonized human glioblastoma single-cell chromatin accessibility atlas (snATAC-seq, 10 studies, 67 patients, 1.65M nuclei, 2.08M peaks)

**DOI:** 10.64898/2026.11.20.768241

**Authors:** Sai Mallannagari, Somto Ngwudike, Jehad Mahmoud, Yanelisse Luiggi Garcia, Victor Carreon, Riya Kajal Rathod, Steven Aguirre, Benjamin Arenkiel, JP McGinnis

## Abstract

As advancing molecular technologies begin to offer exquisite control over cellular processes, the bottleneck becomes our ability to deliver those tools into the specific cell types we want. The paradigm of identifying cell-type-specific enhancer sequences and testing their ability to drive cell-type-specific expression of transgenes has been tried with relatively impressive success in mouse models. It has not been tried to the same degree with enhancer data from humans, and certainly not in human neurological diseases that could benefit from cell-type-selective therapies. Because the first step of this process is compiling single-nucleus ATAC-sequencing data to identify cell-type-specific regions of open chromatin, we present a harmonized atlas of all publicly available IDH-WT glioblastoma snATAC-seq data at cell-type- and patient-level resolution (10 studies, 67 patients, 1,650,376 nuclei total). These are unified into a reference set containing 2,085,175 peaks. 75.7% of distal peaks are absent from the CATLAS reference adult human brain peak set, establishing the novel GBM-specific chromatin landscape across involved cell types. Peaks within recurrent CNV loci (chr7 EGFR, chr12 MDM2, chr8 MYC, chr9 CDKN2A) were flagged to distinguish copy-number inflation from genuine chromatin states. We generated per-cell-type candidate lists—malignant tumor cells, tumor-associated macrophages, resident microglia, astrocytes, oligodendrocytes, OPCs, excitatory neurons, inhibitory neurons, endothelial cells, T cells, and an unassigned but likely-malignant cluster. We focus on peaks ≥ 2 kb from transcriptional start sites and detected in ≥60% of evaluable patients (where a patient is evaluable for a given cell type if they contribute at least 50 quality cells of that type); a peak counts as detected in a patient if ≥1% of that patient’s cells of the target type carry ≥1 Tn5 fragment at the peak. Peaks are then scored per cell type by the mean of two percentile ranks: for strength, the patient-averaged, width-normalized Tn5 fragments per cell at the peak; and for specificity, the log2FC of (target-cell-type strength/next-highest-cell-type strength). Top hits recover expected lineage markers as top-ranked elements in their expected cell types—KDR at rank 1 of the endothelial list, CD3D at rank 4 of the T cell list, and MAG at rank 29 of the oligodendrocyte list—and nominate de novo distal enhancers for every lineage. This atlas is intended to facilitate cell-type-directed therapeutics but will be useful to the broader glioblastoma field. We have made openly available the full ranked lists for all cell types, an interactive and user-friendly HTML viewer, and the underlying pseudobulk matrix, along with per-cell AnnData h5ads for each cohort and an integrated h5ad file encompassing all included studies.

## Introduction

Glioblastoma remains uniformly fatal despite decades of research^1^. Neuroscience more broadly is moving toward interventions—CAR specificity, myeloid reprogramming, vascular normalization, optogenetics, targeted gene editing—all converging on the premise that therapeutic efficacy in the brain requires acting on specific cell types^2^. While cell-type-specific therapies hold great promise for selective manipulation of the brain’s diseases, including glioblastoma, these approaches necessarily rely on molecular features frequently unique to humans and unique to the diseases. If the regulatory state we want to exploit or reprogram is present only in the human tumor microenvironment, looking for it in mouse models or even in normal human brain will fail to recover what could be therapeutically valuable strategies.

Cell-type-selective delivery strategies primarily consist either of modifiable delivery vehicles (such as AAVs) or selective enhancer elements that can switch on transcription only in the desired cell type^3^. The revealed consensus of the neuroscience community is that cell-type-selective tools and therapies are an absolute imperative, with approximately $20M per year over this decade spent on mouse and NHP work by the NIH, including the BRAIN Armamentarium program^4,5,6,7,8,9,10,11,12,13^. Billions more are spent on the industry side. The difficulty is that virtually all this effort is focused on generating technologies highly specific to animal models, which then fail to translate into meaningful therapies for human patients. Two-thirds of regulatory sequences in the human brain do not have mouse correlates^14^. This harmonized, human glioblastoma (WT) snATAC-seq atlas is an effort to identify human regulatory elements that may facilitate such cell-type-selective therapies in glioblastoma, combining all ten publicly available studies that use human patient samples^15-24^.

Enhancer-based cell-type targeting has not been systematically attempted in glioblastoma, though it would allow specific strategies that current cytotoxic and bulk-tumor approaches cannot. Malignant cells could be targeted with cytotoxic, differentiation-forcing, immune-promoting, or anti-migratory payloads. Tumor-associated macrophages could be reprogrammed. Microglia at the infiltrative periphery could be pushed to more aggressively anti-tumor phenotypes. (With the appropriate enhancers, these two related populations could be separately targeted.) T cell targeting could create libraries of CAR-T cells in situ, amidst actively cycling tumor cells. Endothelial cells can be targeted for blood-brain-barrier normalization, or to facilitate therapeutic access. Both reactive astrocytes and neurons could be targeted to reduce bystander toxicity, and reduce excitatory behavior that may influence tumor behavior. A payload under an oligodendrocyte-progenitor enhancer selectively active in the infiltrative edge could address the migratory population that current therapies do not reach. None of these strategies are likely to work using enhancers identified and validated in mouse brains, or even normal human brain. For that, we need data from human glioblastoma samples.

We have therefore compiled a harmonized atlas of all publicly available IDH-WT glioblastoma snATAC-seq data at cell-type- and patient-level resolution (10 studies, 67 patients, 1,650,376 nuclei total). This atlas is intended to facilitate cell-type-directed therapeutics but will be useful to the broader glioblastoma field.

## Results

### Atlas design

We sought to include all publicly available human glioblastoma (IDH-WT) snATAC-seq datasets for which patient- and cell-level peaks were available (this included datasets with available per-cell fragment files and datasets with per-cell peaks annotated but without fragment files). We identified ten datasets, comprising 67 patients, 1.65M nuclei, 2.08M peaks (Table 1, Figure 1).

**Table 1.** Human GBM snATAC-seq cohorts, ordered by data-access tier. All ten included cohorts are open-access. Tier 1 releases raw Tn5 fragments (re-callable against any coordinate system); tier 2 releases per-cell count matrices only (lifted onto the atlas but not re-callable at novel intervals).

| Study | DOI | Accession | Platform | Pts | Nuclei | Frags | Access modality |
|---|---|---|---|---|---|---|---|
| Tier 1 | Open deposit, raw Tn5 fragments released |  |  |  |  |  |  |
| Sojka et al. 2025<br>Nat Cell Biol | 10.1038/s41556-024-01583-9 | GEO GSE276177 | 10x snATAC | 3 | 289,242 | ● | processed + raw fragments |
| Sundaram et al. 2024<br>Science (via TCGA/GDC) | 10.1126/science.adk9217 | GDC blob a7765b5f… | 10x snATAC | 8 | 157,685 | ● | processed + raw fragments |
| Pai et al. 2025<br>Cancer Research | 10.1158/0008-5472.CAN-24-4708 | GEO GSE286415 | 10x snATAC + RNA | 5 | 24,692 | ● | processed + raw fragments |
| Terekhanova et al. 2023<br>Nature (CPTAC-GBM) | 10.1038/s41586-023-06682-5 | GEO GSE240822 | 10x snATAC | 14 | 102,016 | ● | processed + raw fragments |
| Tier 2 | Open deposit, processed per-cell matrices only |  |  |  |  |  |  |
| de Jong et al. 2026<br>Nature (Multiome subset) | Nature 2026 (in press) | AE E-MTAB-17183 | 10x Multiome | 12 | 1,041,869 | ○ | processed h5ad only |
| Lin et al. 2026<br>Nat Neurosci | 10.1038/s41593-026-02265-5 | CNCB HRA004942 | snATAC (low-thru) | 11 | 6,735 | ○ | processed h5ad only |
| Wang X et al. 2024<br>Sci Adv (primary-recurrent) | 10.1126/sciadv.adn4306 | CNGB CNP0003766 | DNBelab C4 snATAC | 5 | 14,378 | ○ | processed h5ad only |
| Wang L et al. 2019<br>Cancer Discovery | 10.1158/2159-8290.CD-19-0329 | GEO GSE138794 | 10x snATAC | 3 | 6,614 | ○ | processed h5ad only |
| Guilhamon et al. 2021<br>eLife (GSC-enriched) | 10.7554/eLife.64090 | GEO GSE139136 | 10x snATAC | 4 | 3,818 | ○ | processed h5ad only |
| Raviram et al. 2023<br>PNAS (sci-ATAC) | 10.1073/pnas.2210991120 | GEO GSE165037 | sci-ATAC | 2 | 3,327 | ○ | processed RDS only |
| Atlas totals (included): 10 cohorts · 67 patients · 1,650,376 nuclei · 2,085,175 union peaks |  |  |  |  |  |  |  |

Frags column: filled green dot = raw per-cell Tn5 fragment files released; hollow dot = processed matrices only.
All deposits are reachable without a data-access-committee application. One additional cohort known to exist at atlas build time was not publicly deposited (Kint, Younes et al. 2025, bioRxiv, 24 patients, Gallo/Fan/Nikolic; not included).
Atlas snapshot v2026-11-10d · Zenodo concept DOI 10.5281/zenodo.23252323 (version DOI 10.5281/zenodo.24149791).

Building a usable atlas from ten independently generated cohorts required decisions that another group could reasonably have made differently. The union set of all peaks was constructed by combining all CATLAS 544K adult-human-brain peaks with novel MACS3 peaks called per pseudobulk (cohort × patient × cell type) on each cohort with fragments available. 75.7% of the resulting 2,085,175 union intervals are de novo relative to CATLAS. Fragments from each cohort were re-scored against this shared peak reference, and cell-type labels were reconciled to a standardized set. In datasets that deposited per-cell peak data already called, without access to fragments, we used the studies’ own peak calls. We did not aggregate adjacent peaks overlapping by less than 100 base pairs, given our interest in identifying shorter cell-type-specific sequences. We focused on individual patient samples as the most important unit of analysis, and so created pseudobulked data for each cell type within each patient’s sample, rather than collapsing each cohort to a single averaged profile. We required at least 50 cells of a given cell type per patient to count towards pseudobulk data; patients with smaller numbers were skipped, hence many cell types have fewer than 67 patients considered evaluable.

IDH-WT glioblastoma almost invariably shows gain of chromosome 7 and loss of chromosome 10, which can artificially inflate accessibility metrics of chromosome 7 and deflate 10 (along with any other copy number variation)—these are dosage artifacts, not real enhancer biology. Ideally this would be controlled in a way specific to each patient, adjusting for their own copy number composition. Since we lack fragment-level for approximately half the patients, we currently just flag (“AMP”) any candidate enhancer sequences found in the following regions prone to copy number variation: chr7p11.2 (EGFR amplification), chr12q15 (MDM2 amplification), chr8q24 (MYC amplification), chr9p21.3 (CDKN2A deletion). This leaves open the option for groups to make their own adjustments. We further flag these regions in non-malignant cells since ambient contamination from these amplified regions was found in other cell type calls, with the amount of contamination correlating with the number of malignant cells found in each dataset.

Tumor cells were identified by glioblastoma chromosomal signature (gain of chromosome 7, loss of chromosome 10), using one of three methods depending on each study’s data (paired single-cell RNA data when available, imputed gene activity from raw sequencing fragments when fragments were released, or a direct chromosome-7-vs-chromosome-10 accessibility ratio when only processed cell-by-peak matrices were deposited). Tumor cells were then further sorted into the four published glioblastoma cellular subtypes (Neftel et al. 2019: astrocyte-like, mesenchymal, neural-progenitor-like, and OPC-like) wherever marker genes could be scored, and each subtype was analyzed as its own population in addition to contributing to a pooled malignant group.

The field of snATAC-seq, especially as it relates to identifying putative enhancer elements for targeted therapies, has not converged on a single metric that predicts enhancer performance. Most seek to balance specificity with strength in some way. Our primary score is a simple average of each peak’s strength rank and its specificity rank. Strength of accessibility is defined as an unweighted per-patient, per-cell-type averaged fragment counts per cell (i.e., if we pull a microglia from patient X, how many fragments at this peak are we expecting to see, averaged across patients X/Y/Z…). Strength is further width-adjusted, to avoid longer sequences looking stronger merely because they have more opportunity for fragment insertion. Specificity is defined as the log2FC of the peak’s strength in that cell type compared to the cell type with the second highest strength. Rankings only evaluate distal elements (>2kb from transcriptional start site) to avoid proximal promoter elements that historically have been poor enhancer candidates.

These choices, the union set of peaks, patients as the replication unit, copy-number flagging instead of adjustment, the primary score equally weighting ranks of specificity and strength, among others, are made explicit so that users wishing to re-score the matrix for their own purposes can easily do so.

### Atlas-level findings

Datasets range from 2 patients with 3,327 nuclei to 12 patients with 1,041,869 nuclei; malignant cells compose 8.8% to 89.9% of sample nuclei, reflecting the infiltrative nature of glioblastoma and the varying proportion of cell types in regions within and adjacent to tumors (Figure 1).

**Figure 1.**
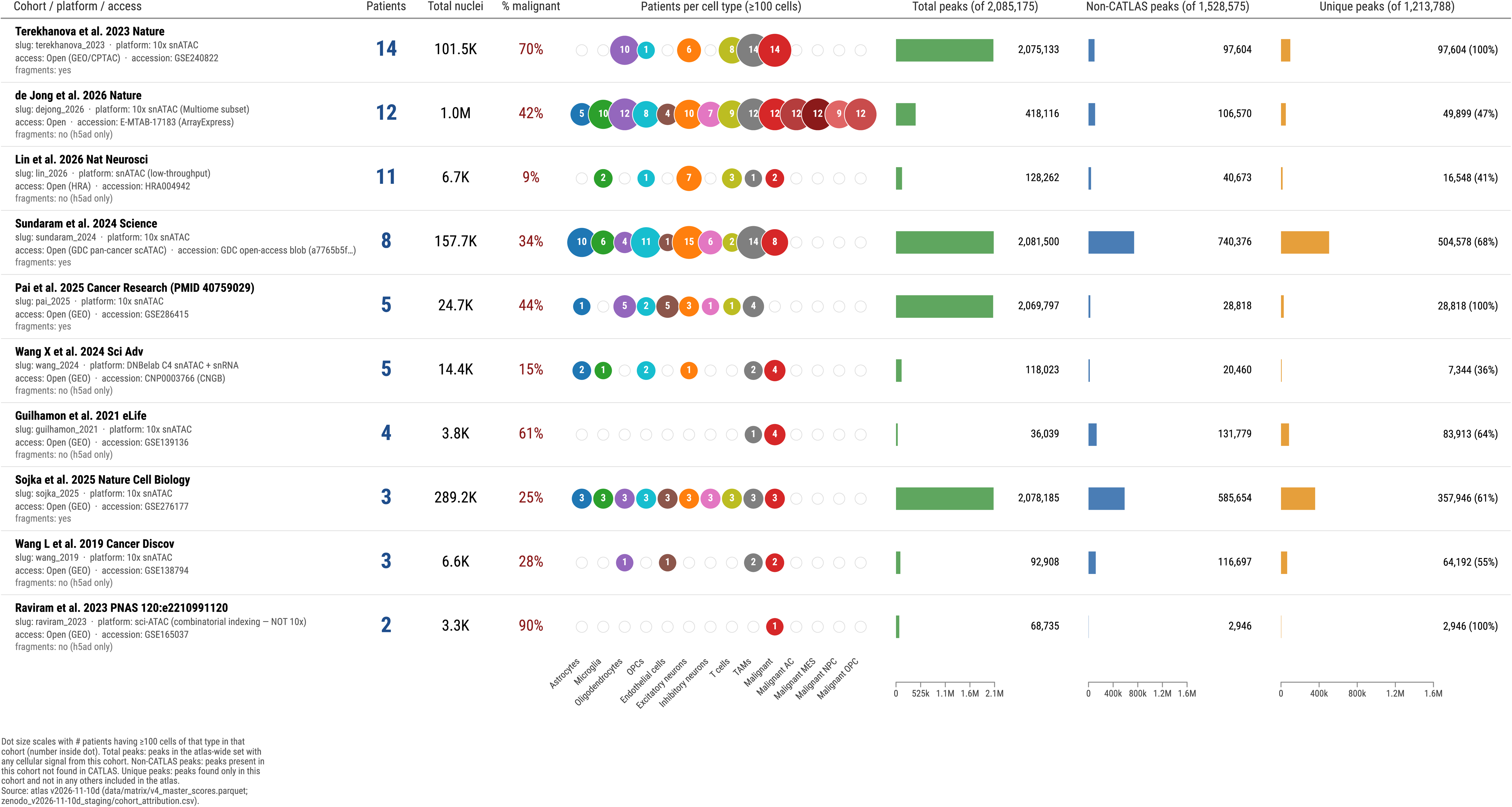
Cohort contributions to the GBM snATAC-seq atlas. Each row describes one of the ten cohorts included in the atlas, ordered chronologically by publication year. Left block: first-author citation, sequencing platform, data-access status and accession. Middle block: patients contributed, total nuclei after cell-level QC, and percentage of cells labelled malignant. Patients-per-cell-type block: filled circles sized by the number of patients in that cohort contributing at least 100 cells for each of the ten primary cell types and four Neftel malignant subtypes used in the atlas. Right block: per-cohort peak contributions, plotted as a share of three atlas-wide denominators — total union peaks (2,085,175), non-CATLAS peaks (1,528,575; atlas peaks absent from the CATLAS normal-brain reference), and uniquely-called peaks (1,213,788; non-CATLAS peaks called in only one cohort). The non-CATLAS and unique bars use a shared scale so the two values are directly comparable within each row. Atlas-wide totals appear in the figure title.

Per-patient nuclei depth varies two orders of magnitude across cohorts (∼600 nuclei per patient in Lin, ∼1k in Guilhamon, ∼2k in Wang L 2019 to ∼96k per patient in Sojka and ∼87k per patient in De Jong 2026). Cell type composition for each study is found in Figure 1. Seven of ten studies used 10X snATAC-seq, one used DNBelab (Wang X 2024), one used sci-ATAC (Raviram), and one used a low-throughput snATAC platform (Lin).

One cohort (De Jong 2026) contributes disproportionately to the atlas: it represents 39% of all pseudobulk columns and has median per-cell-type library size 45−230× larger than the smallest cohorts.

Of the 2.09M peaks, 134,541 are detected in at least 60% of the 67 atlas patients (≥41 of 67 patients, meaning ≥1% of a patient’s cells of the top cell type have a fragment at the defined peak). We use a ‘60% of evaluable patients’ threshold for consistency given that even promoters of canonical housekeeping genes (RPS6, RPL26, POLR2E, GTF2F1, CDC42, SDHB, ribosomal proteins) are recovered in at most 99% of patients (66 of 67, RPS6); the median housekeeping gene has identifiable fragments in just 63% of patients (Supplementary Figure 1A-D). No housekeeping promoter is detected in all 67. Per-peak patient detection is bounded by cross-cohort differences in platform, sequencing depth, and per-donor cell yield. Housekeeping gene promoters further provide a strength (i.e. openness) ceiling of ∼1−2 fragments per cell at their most-accessible cell type (RPL18 1.82, CFL1 1.40, ACTB 0.73) with a median fragments-per-cell of 0.55 (Supplementary Figure 1E). Both per-patient detection and fragments-per-cell recovery of housekeeping genes help calibrate expected values for strong cell-type-specific regions.

78.7% of peaks are distal elements (>2 kb from TSS); 21.3% are promoter-proximal (<2 kb TSS). By genomic location, besides the promoter-proximal regions, 52.2% intragenic (predominantly intronic), and 26.5% intergenic (Supplementary Figure 2A). That promoter-proximal peaks show a substantially higher proportion common to at least 60% of patients (21.3% versus 3.9% of all atlas peaks) demonstrates that promoter accessibility is more universally shared across patients than enhancer accessibility.

CATLAS, the hallmark snATAC-seq human brain atlas, identified 544,735 peaks from their three post-mortem adult donors^14^. We included their entire peak set and combine it with any peak seen in our fragment sets, to generate a union peak set of 2,085,175 total peaks. 75.7% of the peaks in this set are not seen in the CATLAS set (Supplementary Figure 2B). Restricting to peaks that are consistently open in at least one cell type (fragments per cell in their best cell type > 0.1; 70,402 peaks), the CATLAS overlap rises to 87.7% (12.3% novel), indicating that the normal-brain reference captures a large portion of peaks in this atlas, and that most of the novel peaks found in included glioblastoma samples are weaker or less consistent.

When thinking about whether analysis would benefit from further patient samples, the benefit varies by cell type. Overall, the comprehensive peak set is unlikely to meaningfully change, and neurons, astrocytes and OPCs reach their reproducible-peak counts by the sixth chronologically added cohort (Supplementary Figure 2C, 2D). Malignant cells approach but do not fully reach their ceiling. The remaining lineages—TAMs, oligodendrocytes, GABA neurons, endothelial cells, and T cells—continue to be substantially reshaped by each cohort (gaining +38% to +249% of their peak count by the final cohort), largely because one deeply-sequenced study contributes an order of magnitude more cells per lineage than earlier cohorts. Microglia are the clear outlier: the lineage reaches its atlas peak set by cohort 5 but at the lowest absolute peak count of any cell type (∼20k peaks), reflecting the fact that microglia cells are the most sparsely contributed across all 10 cohorts.

Further deeply sequenced patient glioblastoma samples would therefore substantively expand the TAM, oligodendrocyte, and lymphoid/endothelial peak sets, and only refine the nearly saturated neuronal/astrocytic peak sets (Supplementary Figure 2D).

We next asked to what extent peaks cluster into narrow genomic neighborhoods, the pattern commonly associated with super-enhancers. For each cell type we grouped distal peaks by primary cell type (based on counts-per-million) and whose per-cell-type detection rate reached ≥60% of patients, stitching peaks within 12.5 kb of each other. Across the ten primary cell types we identify 17,318 clusters of ≥4 peaks, of which 4,228 contain ≥10 peaks, and 96.6% contain at least one CATLAS peak. The highest-density clusters consistently anchor on lineage-defining or lineage-adjacent genes: ZCCHC24 and DMBT1 for OPCs, EMX2 and GRM3 (PRDM16 ranked #12) for astrocytes, MOBP and MAG for oligodendrocytes, LINC01554, LINC01094 and GPR183 (HMGXB3 ranked #17) for microglia, LAIR1, LILRB1 and SIGLEC10 (CTSB ranked #9) for tumor-associated macrophages, IKZF1 (GZMB ranked #9) for T cells, FOXC1, NOSTRIN and KDR for endothelial cells, VOPP1, LRRN2 and GSE1 for excitatory neurons, and ADARB2-AS1 and CNTN2 (with ADARB2 ranked #20 at the same locus) for GABAergic interneurons (Supplementary Figure 2E).

### Cell-type findings

We restricted cell-type-specific analysis to patient samples containing at least 50 cells of the cell type being analyzed, and to peaks detected in at least 60% of those evaluable patients. For each primary cell type, the number of patients that pass the ≥50-cell floor ranges from 17 (inhibitory neuron) to 58 (malignant), reflecting both the biology of GBM cell-type abundance, surgical specimen retrieval and dissociation, and the sampling depth of each cohort (Figure 2A, 2B). Many of the smaller snATAC-seq datasets do not pass even the 50-malignant-cell threshold due to low cell numbers or sorting for immune cells. Using strength-based primary cell-type assignment, between 2,577 (malignant) and 107,385 (excitatory neuron) reliably open distal peaks are primarily attributable to each cell type at ≥60% patient detection (Figure 3A).

**Figure 2.**
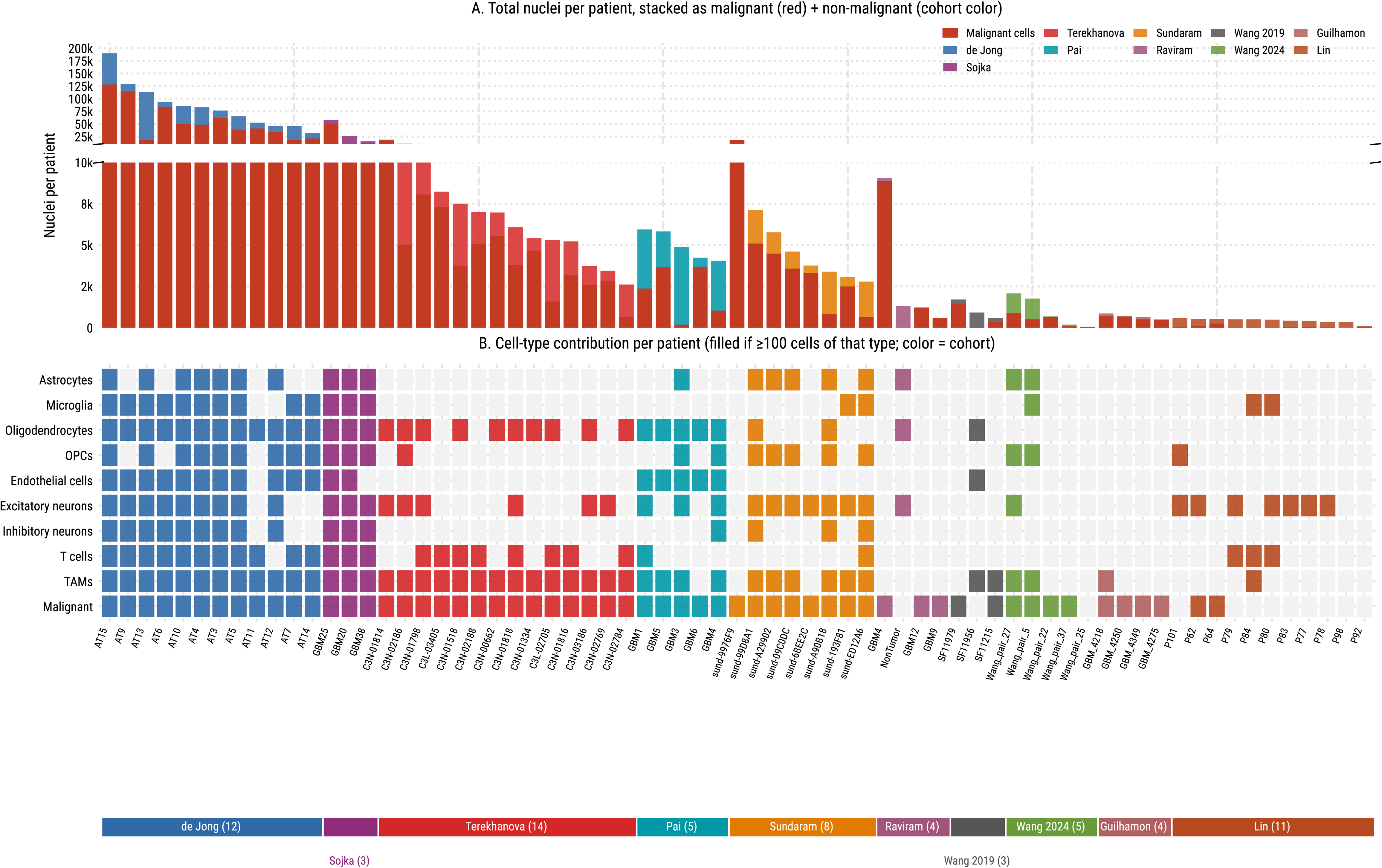
Per-patient cohort composition of the GBM snATAC-seq atlas. Each column is one of the 69 patients in the atlas, ordered by cohort (by cohort median total nuclei, descending) and within cohort by total nuclei. (A) Total nuclei per patient after cell-level QC, stacked as malignant cells (red) and non-malignant cells (cohort color). The y-axis is split at 10,000 nuclei: the upper panel resolves deeply-sequenced patients (10k–200k nuclei) and the lower panel resolves lower-depth patients (0–10k). (B) Cell-type contribution per patient. Rows are the ten primary cell types used in the atlas; a cell is filled (in cohort color) when the patient contributes at least 100 cells of that cell type, and grey otherwise. The strip at the bottom groups patients by cohort and gives each cohort’s patient count in parentheses.

**Figure 3.**
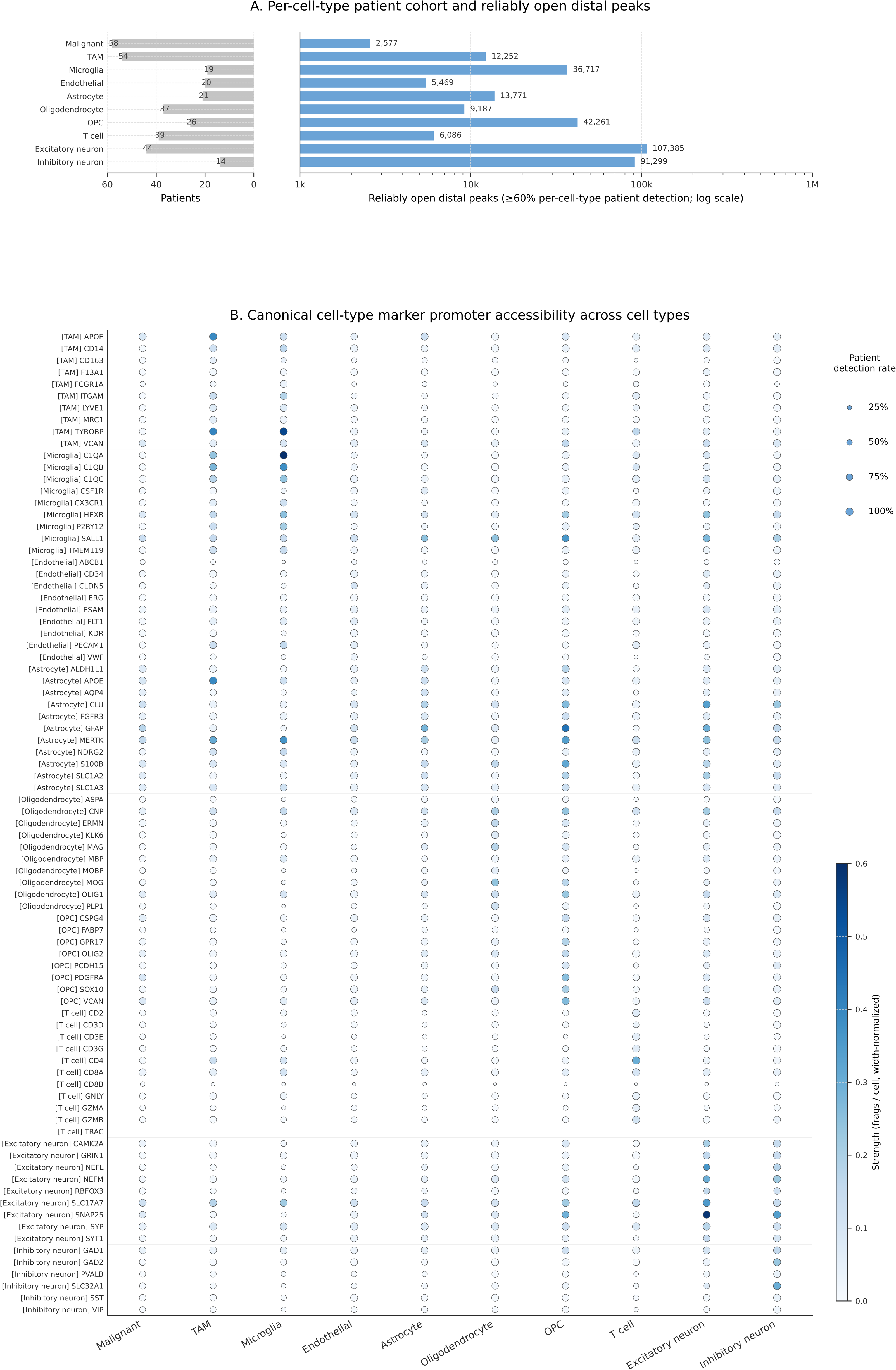
Canonical cell-type markers and the per-cell-type patient / peak inventory. (A) Per-cell-type patient cohort and reliably open distal peaks. Left bars (grey) = number of patients contributing ≥50 cells of each primary cell type to the aggregated atlas. Right bars (light blue, log x-axis) = number of distal peaks reliably open (per-cell-type patient detection ≥ 60%) and primarily attributable to each cell type (strength-based primary; range 2,577 malignant to 107,385 excitatory neurons post-AT9 exclusion). Cell types shown in canonical atlas order. (B) Canonical cell-type marker promoter accessibility. 81 canonical lineage markers (rows, grouped by source cell-type class on the left) × 10 primary cell types (columns). Dot size = patient detection rate; dot color = width-normalized fragments per cell (strength). Promoter peak = highest-strength peak within ±2 kb of the gene’s TSS.

In addition to ubiquitously expressed housekeeping genes, standard cell-type marker genes provide important references for calibrating other atlas findings. Across 67 patients and 1.35 million nuclei spanning ten cohorts, we identified promoter-adjacent peaks (within 2 kb of the TSS) for 81 canonical lineage markers (GFAP, AQP4, SLC1A2, SLC1A3, ALDH1L1, S100B for astrocytes; P2RY12, TMEM119, CX3CR1, CSF1R, C1QA, C1QB, HEXB for microglia; MBP, MOG, PLP1, MAG, MOBP, CNP for oligodendrocytes; and analogous panels for the other six primary cell types with lineage-defining markers) and 12 standard housekeeping genes (ACTB, GAPDH, TUBB, RPL13A, PGK1, TBP, B2M, PPIA, UBC, HMBS, YWHAZ, SDHA). Marker peaks are enriched at their expected cell type but, consistent with GBM biology, often remain substantially accessible in related lineages — excitatory neuron markers (SNAP25, SYT1, NEFL, NEFM, SYP) are shared across both neuronal classes; astrocyte markers (GFAP, SLC1A2, SLC1A3) are also accessible in malignant cells, reflecting the astrocyte-like Neftel states of many GBM tumors; and canonical OPC marker PDGFRA, commonly also used as a malignant-cell marker in GBM, overlaps with both normal OPC and astrocyte programs.

Established marker gene promoters reach a median detection rate of 93% of patients in their expected cell type (range 5−100%) and strength values spanning 0.00 to 0.90 fragments per cell (median 0.12) (Figure 4A). Promoter-level cell-type specificity is modest: only 22 of 81 markers (27%) exceed 2× (log2FC ≥ 1) accessibility in their target cell type compared to the next-most-accessible cell type at the promoter peak, and only 8 of 81 (10%) exceed 4× (log2FC ≥ 2). This follows the well-established finding that cell-type-selective gene regulation is encoded primarily at distal enhancers rather than at promoters^25,26^. Promoters tend to be constitutively accessible across lineages where the gene is expressed, lowly expressed, or poised. For the enhancer-discovery analysis that follows, we therefore constrain our search to distal regulatory elements (≥ 2 kb from any TSS), where chromatin accessibility more faithfully reflects cell-type-selective transcriptional programs.

**Figure 4.**
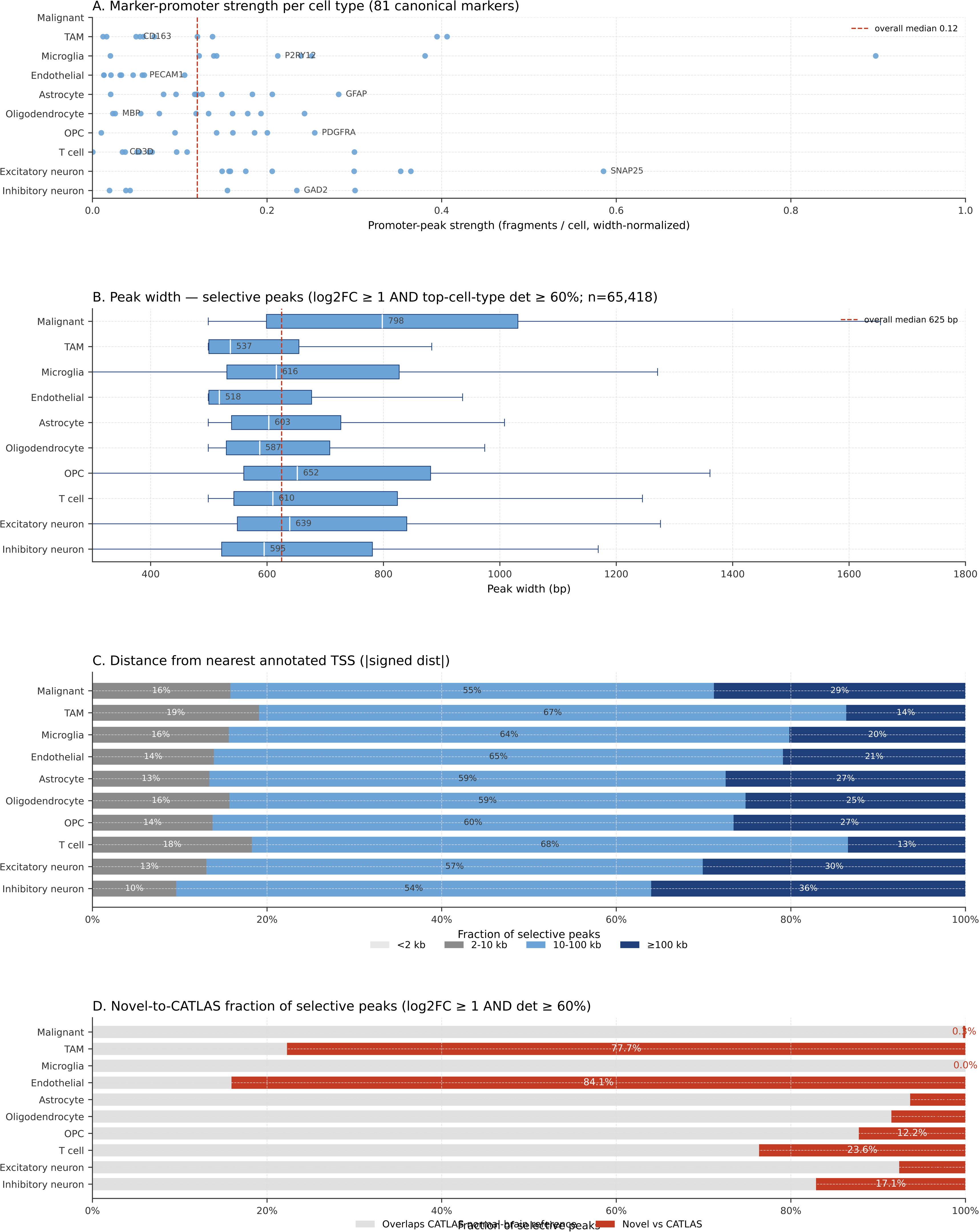
Characterization of cell-type-selective distal peaks (log2FC ≥ 1 vs next-most-accessible cell-type AND top cell-type patient detection ≥ 60%). Across cell types, 65,418 peaks meet the per-cell-type selective criterion. (A) Marker-promoter strength distribution per cell type. One dot per canonical marker at its promoter strength in the marker’s target cell type; red dashed line = overall median across the 81 canonical markers. Illustrative household-name markers labeled per cell type. (B) Peak-width distribution per cell type for the selective peak set. Box = IQR, line = median, whiskers = 1.5× IQR. Per-cell-type median annotated inside each box; red dashed line = overall median across all selective peaks. (C) Distance from nearest annotated TSS. Stacked bars show the fraction of each cell type’s selective peaks in each distance bin: <2 kb, 2–10 kb, 10–100 kb, ≥100 kb. (D) Novel-to-CATLAS fraction. Stacked bars show the fraction of each cell type’s selective peaks overlapping the CATLAS normal-brain reference (grey) or absent from it (red). Per-cell-type novel fractions (post-AT9): Endothelial 84%, TAM 78%, T cell 24%, inhibitory neuron 17%, OPC 12%, oligodendrocyte 9%, excitatory neuron 8%, astrocyte 6%, malignant 0.3%, microglia 0%.

We examined the typical peak width across all peaks replicating in ≥60% of patients and selective for each cell type (log2FC ≥ 1) and found median widths of 500−795 bp per cell type (median across all selective peaks 625 bp, interquartile range 500-780 bp), comfortably within the ∼500 bp AAV cargo window (Figure 4B).

These peaks are overwhelmingly far-distal: 46−65% sit between 10 and 100 kb from any annotated transcription start site, with another 13−39% more than 100 kb from any gene (Figure 4C).

When considering all reliably-open peaks in each cell type (≥60% of patients), novel-to-CATLAS fractions span 0−78% per cell type, with most brain-resident lineages (microglia, astrocyte, oligodendrocyte, OPC, both neuronal classes, malignant) in the 0−15% range and TAM/endothelial as outliers, each above 70%. This separation sharpens when focusing on the cell-type-selective subset for each cell type (log2FC ≥ 1 over the next cell type, detection ≥ 60% of patients). Endothelial (84%) and TAM (78%) selective peaks are overwhelmingly novel regulatory elements absent from normal-brain references, consistent with their tumor involvement that CATLAS does not sample (Figure 4D). T cells (24%) and inhibitory neurons (17%) sit in a middle tier. OPC (12%), oligodendrocyte (9%), excitatory neuron (8%), and astrocyte (6%) selective peaks sit almost entirely within the CATLAS normal-brain regulatory set. In contrast, microglia selective peaks are entirely drawn from the normal-brain repertoire (0% novel; 6,065 selective candidates) and interestingly malignant selective peaks also overlap CATLAS almost completely (0.3% novel; 361 selective candidates).

### Candidate enhancer sequences for cell-type-targeted delivery

The atlas’s primary purpose is to identify regulatory elements that can be packaged into AAV or similar vectors and drive cell-type-selective transgene expression in glioblastoma patients. Candidate enhancers are evaluated across four dimensions. First, strength is defined as ‘if you selected e.g. a microglia at random from any patient in the atlas, how many fragments would you expect to find at this peak’ (per-cell Tn5 fragments at the peak, width-normalized to a 500 bp reference). The more fragments a given cell type has at that peak, the more reliably open the chromatin is in that cell type. (In almost all cases, a cell can at most have only two fragments recovered for any single peak.)

Second, selectivity, which we defined as the log2FC of per-cell accessibility in the target cell type vs. the next-highest cell type. We compare to the second-highest cell type rather than the median or average of all other cell types because median or averages of ‘all others’ can mask a prominent clinically relevant second-highest cell type if all others are quite low. However, reduced specificity may be tolerated depending on the off-target cell types involved. An enhancer sequence that mainly drives expression in malignant cells with off-target expression in neurons is different from a sequence with off-target expression in tumor-associated macrophages, which are generally more dispensable than neurons.

We also highlight patient-level replicability (the fraction of evaluable patients in whom the peak is detected) and cross-cohort replication (the number of distinct cohorts the peak is found in), to guard against candidates that look strong and selective but only in a small number of patients or one cohort. In practice, even strongly expressed housekeeping genes are detected in only ∼87% of (gene × cell-type) patients on average.

Peak coordinates are presented with nearest annotated gene for context only, since enhancer-to-gene regulatory relationships are a separate biological question that AAV-level validation does not necessarily depend on or benefit from. In the sections below, we highlight for each broad cell type representative candidates in four categories, confined to those seen across at least 60% of patients with enough cells to evaluate: canonical lineage anchors, highest-specificity candidates (top log2FC), highest-strength candidates (top fragments per cell), and balanced candidates that score strongly on both specificity and strength. The full ranked candidate set for every cell type is available in the public atlas’s filterable HTML interface and in the supplementary data tables; the candidates we feature here are examples, not an exhaustive shortlist, and users who prefer a different stringency or weighting are encouraged to apply their own filters directly to the atlas.

### Malignant/tumor cells

58 of 67 patients contribute ≥50 malignant cells to the aggregated data; the other samples fall below the floor due either to enrichment protocols or low-input samples. The pan-malignant distal regulatory landscape contains 293,337 peaks reliably open in ≥60% of malignant-evaluable patients, with 2,577 of those most open in malignant cells (Figure 5A). 361 peaks are both reliably identified and selective (log2FC ≥ 1); 9 of those are also strong (≥0.2 frags/cell). Figure 5B traces how the candidate pool shrinks as reliability, selectivity, and strength constraints are combined, along with the percent not seen in the CATLAS reference.

**Figure 5.**
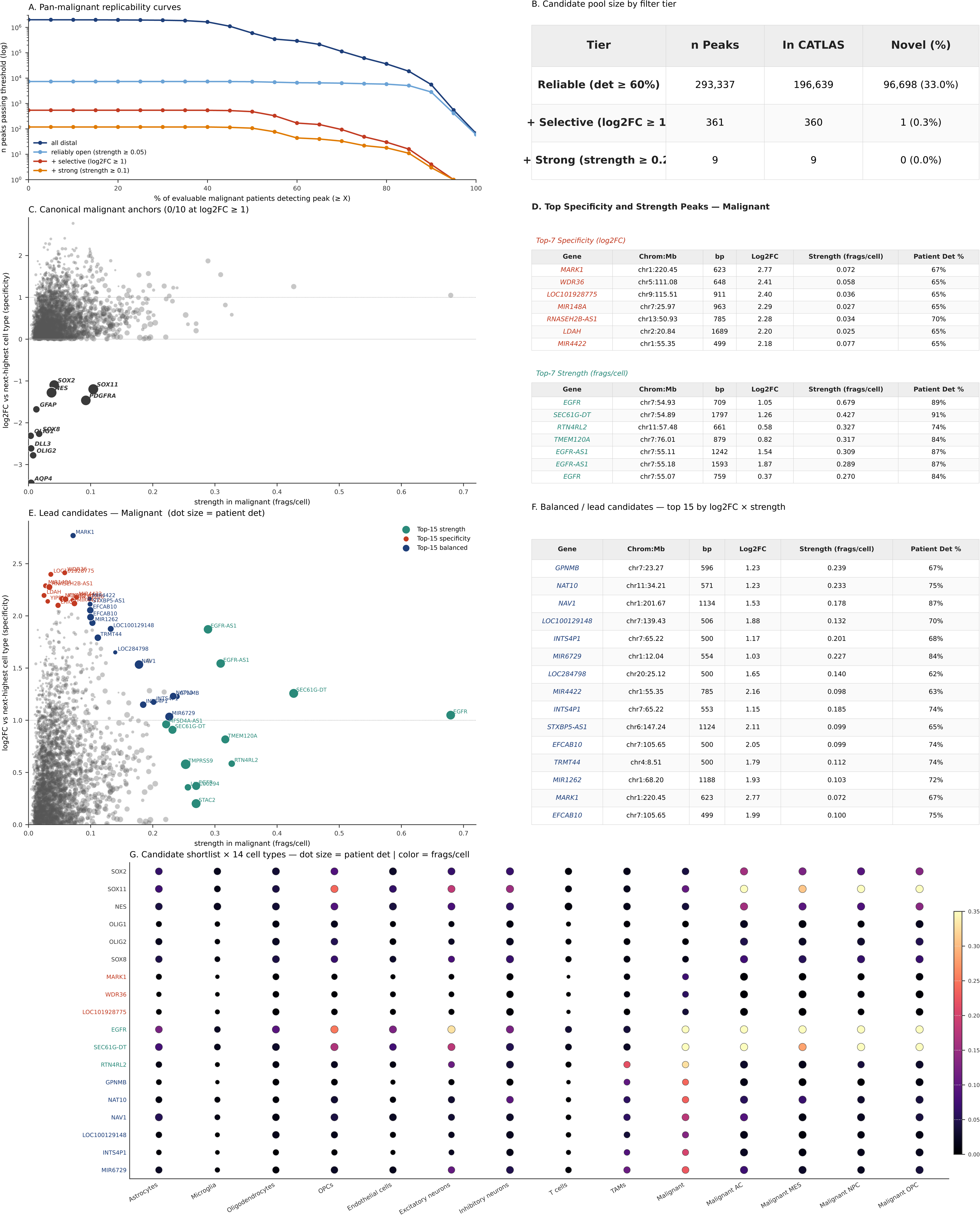
Malignant/tumor-cell candidate enhancers (pan-malignant track, 58 patients with ≥50 malignant cells). (A) Replicability curves — number of distal peaks passing each patient-detection threshold, at four cumulative stringency tiers (all distal / reliably open at strength ≥ 0.05 / + selective at log2FC ≥ 1 / + strong at strength ≥ 0.1); log y-axis. (B) Candidate pool size table — three cumulative stringency tiers (reliable at det ≥ 60%; + selective at log2FC ≥ 1; + strong at strength ≥ 0.2), each with total n, number already in CATLAS, and number novel to CATLAS. (C) Canonical malignant lineage anchors in the strength × specificity space. X = strength in malignant (frags/cell, width-normalized); Y = log2FC vs next-highest primary cell type; dot size = patient detection rate. Charcoal dots = promoter peaks for the 10 canonical anchors (SOX2, SOX11, NES, OLIG1, OLIG2, SOX8, DLL3, AQP4, GFAP, PDGFRA); grey background = all malignant-primary distal peaks at det ≥ 60%. The panel headline reports how many anchors clear log2FC ≥ 1 (0/10 for malignant). (D) Top Specificity and Strength Peaks tables, side by side — Top-7 highest-specificity (log2FC) candidates on the left, Top-7 highest-strength (frags/cell) candidates on the right. Columns: Gene, Chrom:Mb, bp, Log2FC, Strength (frags/cell), Patient Det %. (E) Lead candidates — combined specificity × strength view. X, Y, and dot size as in (C). Top-15 by strength (teal), top-15 by specificity (red), and top-15 by log2FC × strength composite (navy; chr7p11.2 EGFR amplicon locus excluded) are overlaid on the grey background of all malignant-primary distal peaks at det ≥ 60%. Horizontal dotted line = log2FC = 1 selectivity threshold. (F) Balanced / lead candidates — top 15 by log2FC × strength. Columns as in (D). (G) Candidate shortlist × 14 cell types (10 primary + 4 malignant subtypes). Rows = 6 canonical anchors + 3 top-specificity + 3 top-strength + 6 top-balanced candidates. Row label color encodes category (charcoal canonical / red specificity / teal strength / navy balanced). Dot size = patient detection rate; dot color = width-normalized strength.

We find canonical malignant promoter-adjacent peaks across the aggregated cells in 83−98% of malignant-evaluable patients (Figure 5C). These include the stemness transcription factors SOX2, SOX11, and NES; the Neftel-lineage TFs OLIG1, OLIG2, SOX8, and DLL3; the lineage-mimicry markers AQP4, GFAP, PDGFRA, and CD44 and CHI3L1 (mesenchymal); the EGFR locus; and MELK, which is the highest-detection canonical anchor at promoter, open in 98% of malignant-evaluable patients. These canonical anchors cluster at modest log2FC values over the next-highest primary cell type (Figure 5C), reflecting that marker-gene promoters are broadly open across lineages.

The atlas’s cell-type-selective candidates sit substantially above canonical anchors on the specificity axis (Figure 5D, 5E). Among the most specific peaks, we find peaks near MARK1 (log2FC +2.69, or 6.46× the next-most open cell type), WDR36 (log2FC +2.41), LDAH (log2FC +2.20), and DPP10-AS3 (log2FC +2.48), as well as peaks near MIR148A (chr7:25.97 Mb, log2FC +2.29) and LOC101928775 (chr9:115.51 Mb, log2FC +2.18), two uncharacterized regions that nevertheless show both specificity and consistency across patients (Figure 5D, 5E).

Highest-strength candidates (distal peaks, top width-normalized fragments-per-cell at ≥60% patient detection) include EGFR (chr7:54.93 Mb, 90% patient detection, 0.68 frags/cell) and SEC61G-DT (chr7:54.89 Mb, 91% detection, 0.43 frags/cell), both within the chr7p11.2 EGFR amplicon locus whose per-cell strength is partly inflated by local CNV; TMPRSS9 (chr19:2.38 Mb, 97% detection, 0.25 frags/cell); and distal peaks near STAC2 (chr17:39.22 Mb, 91% detection, 0.27 frags/cell), RTN4RL2 (chr11:57.48 Mb, 74% detection, 0.33 frags/cell), and TMEM120A (chr7:76.01 Mb, 85% detection, 0.32 frags/cell) (Figure 5D, 5E).

Balanced candidates (strong on both specificity and strength) include GPNMB (chr7:23.27 Mb, log2FC +1.23, 0.24 frags/cell, 67% patient detection), NAT10 (chr11:34.21 Mb, log2FC +1.23, 0.23 frags/cell, 76% detection), and NAV1 (chr1:201.66 Mb, log2FC +1.53, 0.18 frags/cell, 88% detection).

### Myeloid (combined)

The myeloid compartment in glioblastoma comprises both microglia (brain-resident, yolk-sac-derived) and tumor-associated macrophages (TAMs, recruited from circulating monocytes), which share substantial chromatin accessibility profiles. Eight of ten cohorts explicitly separated microglia from TAM at the cell-label level; one labels all myeloid cells as TAMs without a dedicated microglia class. We generated scores for three myeloid groupings: a pan-myeloid set (microglia + TAM combined, 58 of 67 patients contributing ≥50 myeloid cells), a microglia-only set (30 patients at ≥50), and a TAM-only set (54 patients at ≥50 cells). The pan-myeloid distal regulatory landscape contains 282,773 peaks reliably open in ≥60% of myeloid-evaluable patients; of these, 477 are both selective (log2FC ≥ 1) and strong (≥0.2 frags/cell) (Figure 6A, 6B). The strength and specificity of regions near canonical myeloid markers are shown relative to other regions in Figure 6C1−C3.

**Figure 6.**
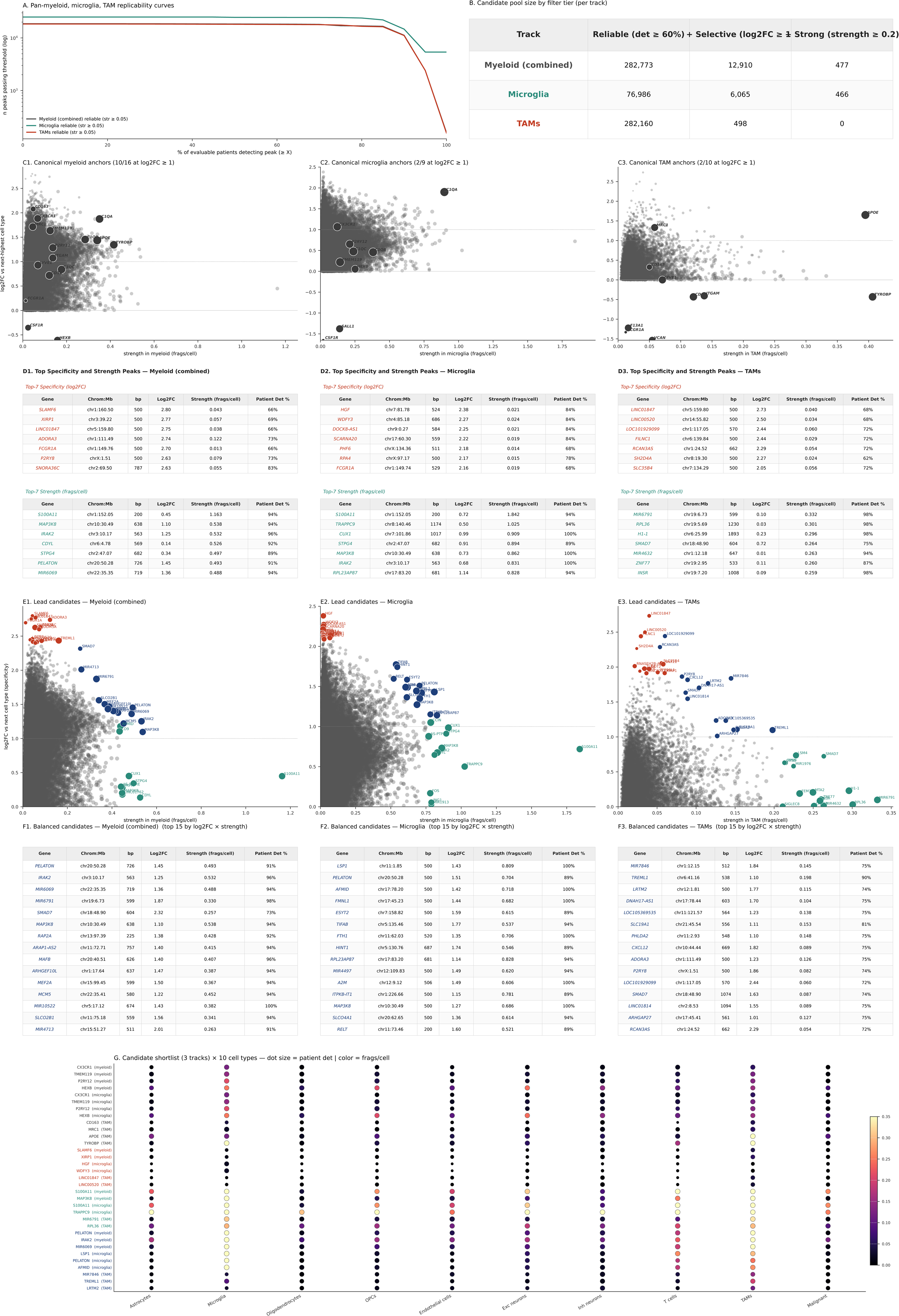
Myeloid (pan-myeloid, microglia-only, TAM-only) candidate enhancers. Three tracks: Myeloid (combined), Microglia, TAMs. 58 of 67 patients contribute ≥50 myeloid cells; 30 patients ≥50 microglia; 54 patients ≥50 TAM cells. (A) Replicability curves — number of distal peaks reliably open (strength ≥ 0.05) in each track, as a function of patient-detection threshold. Three curves overlaid (grey myeloid, teal microglia, red TAM); log y-axis. (B) Candidate pool size table — three tracks × three cumulative stringency tiers (reliable, + selective, + strong). (C1–C3) Canonical myeloid anchors in the strength × specificity space, one scatter per track. Myeloid (combined) scored against other cell types excluding both microglia and TAMs so canonical myeloid markers show true specificity (10/16 at log2FC ≥ 1). Microglia panel uses the microglia canonical set (2/9 at log2FC ≥ 1); TAM panel uses the TAM canonical set (2/10 at log2FC ≥ 1). Charcoal = canonical anchors; grey = all primary-track distal peaks at det ≥ 60%; dot size = patient detection rate. (D1–D3) Per-track Top Specificity and Strength Peaks — Top-7 specificity + Top-7 strength tables stacked per track. (e1–e3) Per-track lead candidates — combined strength × log2FC scatters overlaying top-15 strength (teal), specificity (red), and balanced (navy) candidate sets on the per-track grey background. (F1–F3) Per-track balanced top-15 tables — gene, Chrom:Mb, bp, log2FC, strength, patient detection. (G) Combined candidate shortlist dotplot — 4 canonical + 2 top-specificity + 2 top-strength + 3 top-balanced per track × 10 primary cell types. Row labels carry the originating track in parentheses; label color encodes category (charcoal canonical / red specificity / teal strength / navy balanced). Dot size = patient detection rate; color = width-normalized strength.

Among the most pan-myeloid-specific peaks are LINC01847 (log2FC +2.75) and XIRP1 (+2.71) (Figure 6E1, 6D1), but these sit at ∼0.04−0.06 frags/cell and are unlikely to drive robust transgene expression. The highest-strength peaks (S100A11 at 1.16 frags/cell, ZNF77 at 0.75) (Figure 6E1, 6D1) have strongly negative log2FC (−0.76 and −0.90) and are broadly accessible outside the myeloid compartment, so not usable as selective sequences.

The balanced pan-myeloid candidates (Figure 6F1, 6G, ordered by log2FC × strength) cluster at log2FC 1.0−2.1 and strength 0.2−0.4 frags/cell and are the lead candidates for pan-myeloid targeting. The top ten include CTSB (chr8:11.90 Mb, log2FC +2.07, 0.21 frags/cell, 73% det)—cathepsin B, the classic myeloid lysosomal protease; CSF2RA (chrX:1.24 Mb, +2.11, 0.19, 73%)—the GM-CSF receptor a subunit, a canonical myeloid-lineage marker; MAF (chr16:79.54 Mb, +1.64, 0.20, 91%); LGMN (chr14:92.66 Mb, +1.21, 0.27, 95%); ADORA3 (chr1:111.49 Mb, +1.24, 0.27, 95%); and SMAD7 (chr18:48.90 Mb, +1.98, 0.26, 73%). MIR6791 (chr19:6.73 Mb, +1.36, 0.33, 98%), ARAP1-AS2 (chr11:72.71 Mb, +1.06, 0.41, 95%), MIR4713 (chr15:51.27 Mb, +1.64, 0.26, 91%), and PRRT1B (chr9:131.60 Mb, +1.01, 0.33, 95%) are additional high-scoring candidates. These would be useful in cases where targeting the entire myeloid compartment is desirable or tolerable.

### Microglia

19 of 67 patients contribute ≥50 microglia cells to the aggregated data, reflecting both microglia depletion during tumor-cell enrichment in some cohorts. The pan-microglia distal regulatory landscape contains 76,986 peaks reliably open in ≥60% of microglia-evaluable patients, of which 6,065 are both reliable and selective (log2FC ≥ 1 over the next cell type) (Figure 6B). The second-highest cell type at microglia-selective peaks is TAMs (75.7% of 6,065 selective peaks), reflecting shared myeloid programs. Canonical marker promoters CX3CR1, TMEM119, P2RY12, HEXB, CSF1R, AIF1, and the C1Q cluster (C1QA, C1QB, C1QC) are all microglia-primary at promoter, but none clear the 2× specificity threshold (log2FC ≥ 1): CX3CR1 at log2FC +0.70 (95% det, 0.31 frags/cell) and C1QA at +0.57 (100% det, 0.33 frags/cell) are the two highest-specificity canonical anchors (Figure 6C2).

Among the most specific microglia distal peaks are DKK2 (log2FC +2.02) and FBXO11 (+1.95) (Figure 6E2, 6D2), but these sit at ∼0.02 frags/cell and are unlikely to drive robust transgene expression. The highest-strength peaks (S100A11 at 1.84 frags/cell, TRAPPC9 at 1.02) (Figure 6E2, 6D2) have log2FC near zero and are not microglia-selective.

The balanced microglia candidates (Figure 6F2, 6G, ordered by log2FC × strength) cluster at log2FC 1.0− 1.4 and strength 0.5−0.8 frags/cell. These are the most promising targeting elements. They include LSP1 (chr11:1.85 Mb, log2FC +1.08, 0.81 frags/cell, 100% det)—the leukocyte-specific protein, already a classical myeloid marker; MAP3K8 (chr10:30.49 Mb, +1.27, 0.69 frags/cell, 100% det)—the inflammatory kinase TPL2, broadly expressed in activated myeloid cells; FTH1 (chr11:62.02 Mb, +1.13, 0.71 frags/cell, 100% det)—ferritin heavy chain; FMNL1 (chr17:45.23 Mb, +1.39, 0.68 frags/cell, 100% det); AFMID (chr17:78.20 Mb, +1.31, 0.72 frags/cell, 100% det); and A2M (chr12:9.12 Mb, +1.20, 0.61 frags/cell, 100% det). RPL23AP87 (chr17:83.20 Mb, +1.14, 0.83 frags/cell, 95% det), PELATON (chr20:50.28 Mb, +1.24, 0.70 frags/cell, 89% det), MIR4497 (chr12:109.83 Mb, +1.31, 0.62 frags/cell, 95% det), and DPYSL2 (chr8:26.57 Mb, +1.25, 0.61 frags/cell, 100% det) are also promising candidates.

### Tumor-associated macrophages

54 of 67 patients contribute ≥50 cells labeled as TAMs. The pan-TAM distal regulatory landscape contains 282,160 peaks reliably open in ≥60% of TAM-evaluable patients, of which 498 are also selective at log2FC ≥ 1 vs the next cell type (Figure 6B). Canonical TAM markers APOE (log2FC −0.94, 94% det, 0.17 frags/cell), SPP1 (−0.75, 96% det, 0.17 frags/cell), MRC1 (−0.71, 87% det, 0.13 frags/cell), CD163 (−0.58, 91% det, 0.14 frags/cell), and LYVE1 (−0.49, 91% det, 0.16 frags/cell) are detected at high patient reproducibility but are all stronger in microglia or malignant cells at the promoter peak, so none reach 2× TAM specificity (Figure 6C2). The second-highest cell type at TAM-selective peaks is typically microglia (55.4% of 498 selective peaks) or neurons.

Among the most specific TAM distal peaks are LINC01847 (log2FC +2.73), PRR16 (+2.30), and TLR4 (+1.98) (Figure 6E3, 6D3), but these sit at only 0.03−0.05 frags/cell and are unlikely to drive robust transgene expression. The highest-strength TAM-primary peaks (S100A11 at 1.12 frags/cell, ZNF77 at 0.76) (Figure 6E3, 6D3) have strongly negative log2FC (−0.81 and −0.88) — they are open at comparable strength in microglia and other lineages and are not TAM-selective.

The balanced TAM candidates (Figure 6F3, 6G, top 15 by log2FC × strength) are notably weaker than the microglia or pan-myeloid balanced sets. The TAM balanced pool contains only 79 candidates total vs ∼1,000 for microglia and pan-myeloid, and top strengths plateau at ∼0.2 frags/cell rather than 0.5−0.8. The top ten include TREML1 (chr6:41.16 Mb, log2FC +1.10, 0.20 frags/cell, 91% det) — triggering receptor expressed on myeloid cells-like 1, a classical myeloid surface receptor; CXCL12 (chr10:44.44 Mb, +1.82, 0.09, 76% det); and ADORA3 (chr1:111.49 Mb, +1.23, 0.13, 76% det). Others are MIR7846 (chr1:12.15 Mb, +1.57, 0.14, 76%), SLC19A1 (chr21:45.54 Mb, +1.11, 0.15, 81%), LRTM2 (chr12:1.81 Mb, +1.39, 0.12, 74%), P2RY8 (chrX:1.51 Mb, +1.86, 0.08, 74%), DNAH17-AS1 (chr17:78.44 Mb, +1.35, 0.10, 76%), ARHGAP27 (chr17:45.41 Mb, +1.01, 0.13, 76%), and EDDM3A (chr14:20.73 Mb, +1.59, 0.08, 76%). Overall, TAM-targeting may be best accomplished via pan-myeloid candidates (Figure 6E1, 6F1), which are both stronger and more cohort-replicable, rather than TAM-exclusive distal elements., so long as that is compatible with the therapeutic strategy.

### Endothelial cells

20 of 67 patients contribute ≥50 endothelial cells to the aggregated data. Endothelial cells are a smallraction of GBM dissociations and often lost in enrichment. The pan-endothelial distal regulatory landscape\ contains 193,368 peaks reliably open in ≥60% of endothelial-evaluable patients, of which 5,469 are most open in endothelial and 1,086 are both reliable and selective (log2FC ≥ 1 vs the next cell type) (Figure 7A, 7B). VWF (+1.23, 75% det) and ABCB1 (+1.76, 40% det) have promoters that clear the 2× specificity threshold at the promoter (Figure 7C).

**Figure 7.**
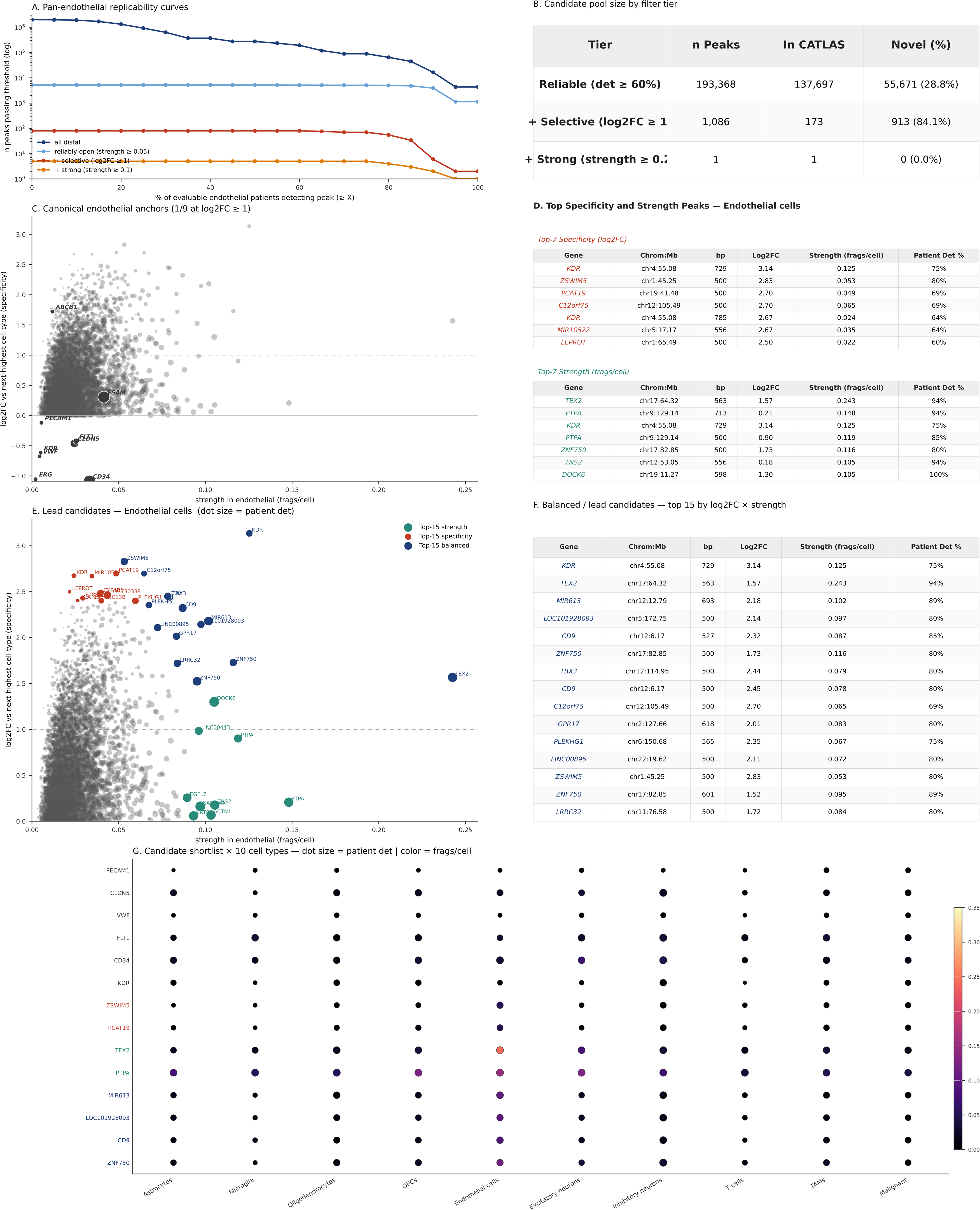
Endothelial cell candidate enhancers (20 patients with ≥50 endothelial cells). (A) Replicability curves — number of distal peaks passing each patient-detection threshold, at four cumulative stringency tiers (all distal / reliably open at strength ≥ 0.05 / + selective at log2FC ≥ 1 / + strong at strength ≥ 0.1); log y-axis. (B) Candidate pool size table — three cumulative stringency tiers (reliable at det ≥ 60%; + selective at log2FC ≥ 1; + strong at strength ≥ 0.2), each with total n, number already in CATLAS, and number novel to CATLAS. (C) Canonical endothelial lineage anchors in the strength × specificity space. X = strength in endothelial (frags/cell, width-normalized); Y = log2FC vs next-highest primary cell type; dot size = patient detection rate. Charcoal dots = promoter peaks for the 9 canonical anchors (PECAM1, CLDN5, VWF, FLT1, CD34, KDR, ESAM, ERG, ABCB1); grey background = all endothelial-primary distal peaks at det ≥ 60%. The panel headline reports how many anchors clear log2FC ≥ 1 (2/9 clear log2FC ≥ 1 for endothelial). (D) Top Specificity and Strength Peaks tables, side by side — Top-7 highest-specificity (log2FC) candidates on the left, Top-7 highest-strength (frags/cell) candidates on the right. Columns: Gene, Chrom:Mb, bp, Log2FC, Strength (frags/cell), Patient Det %. (E) Lead candidates — combined specificity × strength view. X, Y, and dot size as in (C). Top-15 by strength (teal), top-15 by specificity (red), and top-15 by log2FC × strength composite (navy) are overlaid on the grey background of all endothelial-primary distal peaks at det ≥ 60%. Horizontal dotted line = log2FC = 1 selectivity threshold. (F) Balanced / lead candidates — top 15 by log2FC × strength. Columns as in (D). (G) Candidate shortlist × 10 primary cell types. Rows = 6 canonical anchors + 3 top-specificity + 3 top-strength + 6 top-balanced candidates for endothelial. Row label color encodes category (charcoal canonical / red specificity / teal strength / navy balanced). Dot size = patient detection rate; dot color = width-normalized strength.

Among the most specific endothelial distal peaks are KDR (chr4:55.08 Mb, log2FC +2.74, 75% det, 0.13 frags/cell) and TBX3 (chr12:114.66 Mb, +2.33, 75% det, 0.05 frags/cell) (Figure 7D, 7E), but most top-specificity regions cluster at ∼0.03−0.05 frags/cell and are unlikely to drive strong transgene expression (Figure 7D, 7E). The highest-strength endothelial peaks are headed by TEX2 (chr17:64.32 Mb, 0.24 frags/cell, log2FC +1.23, 95% det) and KDR (0.13, +2.74), followed by ZNF750 (chr17:82.84 Mb, 0.12, +1.26), DOCK6 (chr19:11.27 Mb, 0.11, +0.43), and MIR613 (chr12:12.79 Mb, 0.10, +1.40) (Figure 7D, 7E).

The balanced endothelial candidates (Figure 7E, 7F, log2FC × strength) overlap substantially with the top-strength list. The top ten include KDR (chr4:55.08 Mb, log2FC +2.74, 0.13 frags/cell, 75% det), vascular endothelial growth factor receptor 2 (VEGFR2), the canonical endothelial receptor and a well-established vascular target; TEX2 (chr17:64.32 Mb, +1.23, 0.24 frags/cell, 95% det); TBX3 (chr12:114.95 Mb, +2.22, 0.08 frags/cell, 80% det); CD9 (chr12:6.17 Mb, +1.65, 0.09 frags/cell, 85% det); MIR613 (chr12:12.79 Mb, +1.40, 0.10 frags/cell, 90% det); DUSP5 (chr10:110.43 Mb, +1.96, 0.06 frags/cell, 65% det); and PLEKHG1 (chr6:150.68 Mb, +2.00, 0.07 frags/cell, 75% det). ZNF750 (chr17:82.84 Mb, +1.26, 0.12, 80%), LOC101928093 (chr5:172.75 Mb, +1.68, 0.10, 80%), and C12orf75 (chr12:105.49 Mb, +2.11, 0.06, 70%) lack more obvious endothelial connections but score highly (Figure 7F).

### Astrocytes

21 of 67 patients contribute ≥50 astrocyte cells to the aggregated data. The pan-astrocyte distal regulatory landscape contains 174,848 peaks reliably open in ≥60% of astrocyte-evaluable patients, of which 13,771 are most open in astrocytes and 1,642 are both reliable and selective (log2FC ≥ 1) (Figures 8A, 8B).

Canonical marker promoters remain the least specific of any cell type examined: zero classic astrocyte markers clear 2× specificity at the promoter peak. GFAP (−1.58), ALDH1L1 (−0.76), SLC1A3 (−1.20), SLC1A2 (−1.79), S100B (−1.98), FGFR3 (−1.11), NDRG2 (−2.09), CLU (−1.54), MERTK (−1.80), ATP1A2 (−1.23), and APOE (−1.65) are all more accessible at the promoter peak in neurons, TAMs, or OPCs than in astrocytes themselves; only AQP4 (+0.58) and GJA1 (+0.99) are astrocyte-primary at the promoter at all (Figure 8C).

**Figure 8.**
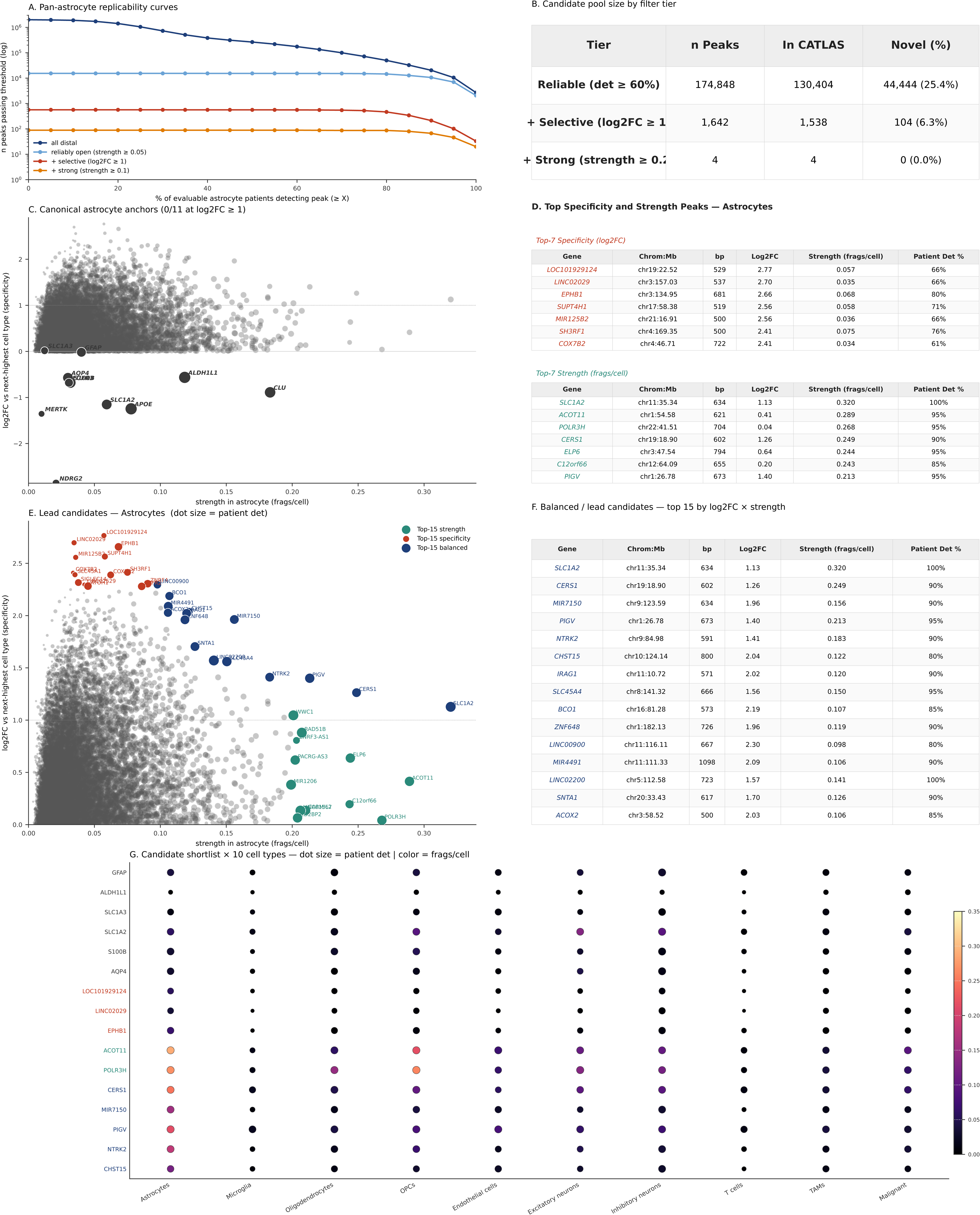
Astrocyte candidate enhancers (21 patients with ≥50 astrocyte cells). (A) Replicability curves — number of distal peaks passing each patient-detection threshold, at four cumulative stringency tiers (all distal / reliably open at strength ≥ 0.05 / + selective at log2FC ≥ 1 / + strong at strength ≥ 0.1); log y-axis. (B) Candidate pool size table — three cumulative stringency tiers (reliable at det ≥ 60%; + selective at log2FC ≥ 1; + strong at strength ≥ 0.2), each with total n, number already in CATLAS, and number novel to CATLAS. (C) Canonical astrocyte lineage anchors in the strength × specificity space. X = strength in astrocyte (frags/cell, width-normalized); Y = log2FC vs next-highest primary cell type; dot size = patient detection rate. Charcoal dots = promoter peaks for the 11 canonical anchors (GFAP, ALDH1L1, SLC1A3, SLC1A2, S100B, AQP4, APOE, CLU, FGFR3, NDRG2, MERTK); grey background = all astrocyte-primary distal peaks at det ≥ 60%. The panel headline reports how many anchors clear log2FC ≥ 1 (0/11 clear log2FC ≥ 1 for astrocyte). (D) Top Specificity and Strength Peaks tables, side by side — Top-7 highest-specificity (log2FC) candidates on the left, Top-7 highest-strength (frags/cell) candidates on the right. Columns: Gene, Chrom:Mb, bp, Log2FC, Strength (frags/cell), Patient Det %. (E) Lead candidates — combined specificity × strength view. X, Y, and dot size as in (C). Top-15 by strength (teal), top-15 by specificity (red), and top-15 by log2FC × strength composite (navy) are overlaid on the grey background of all astrocyte-primary distal peaks at det ≥ 60%. Horizontal dotted line = log2FC = 1 selectivity threshold. (F) Balanced / lead candidates — top 15 by log2FC × strength. Columns as in (D). (G) Candidate shortlist × 10 primary cell types. Rows = 6 canonical anchors + 3 top-specificity + 3 top-strength + 6 top-balanced candidates for astrocyte. Row label color encodes category (charcoal canonical / red specificity / teal strength / navy balanced). Dot size = patient detection rate; dot color = width-normalized strength.

Among the most specific astrocyte distal peaks are LOC101929124 (chr19:22.52 Mb, log2FC +2.77, 67% det, 0.06 frags/cell), LINC02029 (chr3:157.03 Mb, +2.67, 67% det, 0.04), EPHB1 (chr3:134.95 Mb, +2.49, 81% det, 0.07), and BCO1 (chr16:81.28 Mb, +2.21, 86% det, 0.11) (Figure 8D, 8E), but the top-specificity peaks cluster at 0.03−0.07 frags/cell. The highest-strength astrocyte peaks are headed by a SLC1A2 distal element (chr11:35.34 Mb, 0.32 frags/cell, log2FC +0.63, 100% det), ACOT11 (chr1:54.58 Mb, 0.29, +0.41, 95% det), POLR3H (chr22:41.51 Mb, 0.27, +0.04, 95% det), CERS1 (chr19:18.90 Mb, 0.25, +0.76, 90% det), and ELP6 (chr3:47.54 Mb, 0.24, +0.17, 95% det), broadly open across glial cell types and falling below 2× specificity (Figure 8D, 8E).

The balanced astrocyte candidates (Figure 8E, 8F, log2FC × strength) include MIR7150 (chr9:123.59 Mb, log2FC +1.96, 0.16 frags/cell, 90% det); IRAG1 (chr11:10.72 Mb, +2.02, 0.12, 90% det); CHST15 (chr10:124.14 Mb, +1.95, 0.12, 81% det); BCO1 (chr16:81.28 Mb, +2.21, 0.11, 86% det); ZNF648 (chr1:182.13 Mb, +1.96, 0.1, 90% det); LINC02200 (chr5:112.58 Mb, +1.57, 0.14, 100% det); SNTA1 (chr20:33.43 Mb, +1.69, 0.13, 90% det); SLC45A4 (chr8:141.32 Mb, +1.37, 0.15, 95% det); MFGE8 (chr15:88.95 Mb, +1.23, 0.16, 100% det); and ACTL7B (chr9:108.72 Mb, +1.99, 0.10, 86% det). NUDT2, PIK3C2G, LINC02650, and KCNK16 round out the balanced top-15 with scores in the same range but with less obvious astrocyte biology involvement. The second-highest cell type for nearly all astrocyte-selective candidates is malignant_AC, which remains a consideration for astrocyte-targeted therapies in a GBM context.

### Oligodendrocytes

37 of 67 patients contribute ≥50 oligodendrocyte cells to the analysis. The pan-oligodendrocyte distal regulatory landscape contains 319,721 peaks reliably open in ≥60% of oligodendrocyte-evaluable patients, of which 9,187 are most open in oligodendrocytes and 790 are both reliable and selective (log2FC ≥ 1 vs the next cell type) (Figure 9A, 9B). Canonical oligodendrocyte marker promoters are the strongest and most specific of any glial cell type: PLP1 (+2.12, 100% det) clears the 2× specificity threshold, with OPALIN (+1.25), KLK6 (+1.15), MOBP (+0.94), NKX6-2 (+0.86), MAG (+0.85), ERMN (+0.74), MOG (+0.50), and ASPA (+0.41) all remaining oligodendrocyte-primary at the promoter peak with substantial specificity (Figure 9C). The second-highest cell type at oligodendrocyte-selective distal peaks is overwhelmingly OPC (84.4% of the 790 selective peaks), reflecting shared myelin-lineage regulatory programs that may complicate strictly oligodendrocyte-specific transgene targeting in glioblastoma patients.

**Figure 9.**
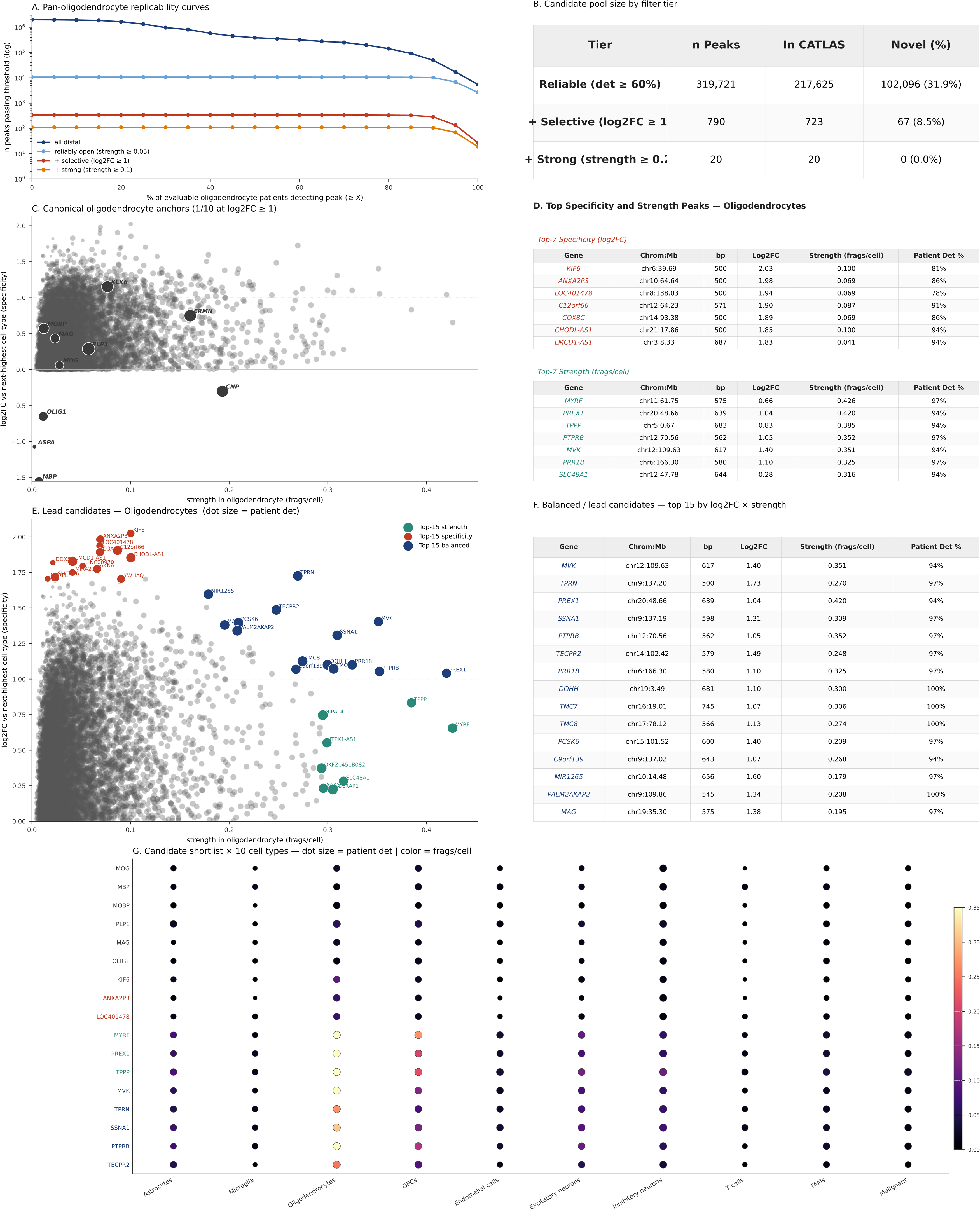
Oligodendrocyte candidate enhancers (37 patients with ≥50 oligodendrocyte cells). (A) Replicability curves — number of distal peaks passing each patient-detection threshold, at four cumulative stringency tiers (all distal / reliably open at strength ≥ 0.05 / + selective at log2FC ≥ 1 / + strong at strength ≥ 0.1); log y-axis. (B) Candidate pool size table — three cumulative stringency tiers (reliable at det ≥ 60%; + selective at log2FC ≥ 1; + strong at strength ≥ 0.2), each with total n, number already in CATLAS, and number novel to CATLAS. (C) Canonical oligodendrocyte lineage anchors in the strength × specificity space. X = strength in oligodendrocyte (frags/cell, width-normalized); Y = log2FC vs next-highest primary cell type; dot size = patient detection rate. Charcoal dots = promoter peaks for the 10 canonical anchors (MOG, MBP, MOBP, PLP1, MAG, OLIG1, CNP, ERMN, KLK6, ASPA); grey background = all oligodendrocyte-primary distal peaks at det ≥ 60%. The panel headline reports how many anchors clear log2FC ≥ 1 (2/10 clear log2FC ≥ 1 for oligodendrocyte). (D) Top Specificity and Strength Peaks tables, side by side — Top-7 highest-specificity (log2FC) candidates on the left, Top-7 highest-strength (frags/cell) candidates on the right. Columns: Gene, Chrom:Mb, bp, Log2FC, Strength (frags/cell), Patient Det %. (E) Lead candidates — combined specificity × strength view. X, Y, and dot size as in (C). Top-15 by strength (teal), top-15 by specificity (red), and top-15 by log2FC × strength composite (navy) are overlaid on the grey background of all oligodendrocyte-primary distal peaks at det ≥ 60%. Horizontal dotted line = log2FC = 1 selectivity threshold. (F) Balanced / lead candidates — top 15 by log2FC × strength. Columns as in (D). (G) Candidate shortlist × 10 primary cell types. Rows = 6 canonical anchors + 3 top-specificity + 3 top-strength + 6 top-balanced candidates for oligodendrocyte. Row label color encodes category (charcoal canonical / red specificity / teal strength / navy balanced). Dot size = patient detection rate; dot color = width-normalized strength.

Among the most specific oligodendrocyte distal peaks are KIF6 (chr6:39.69 Mb, log2FC +2.03, 81% det, 0.10 frags/cell), ANXA2P3 (chr10:64.64 Mb, +1.98, 86% det, 0.07), C12orf66 (chr12:64.23 Mb, +1.90, 92% det, 0.09), COX8C (chr14:93.38 Mb, +1.89, 86% det, 0.07), and CHODL-AS1 (chr21:17.86 Mb, +1.85, 95% det, 0.10) (Figure 9D, 9E). The highest-strength oligodendrocyte peaks are headed by a MYRF distal element (chr11:61.75 Mb, 0.43 frags/cell, log2FC +0.20, 97% det), PREX1 (chr20:48.66 Mb, 0.42, +1.04, 95% det), TPPP (chr5:0.67 Mb, 0.39, +0.83, 95% det), PTPRB (chr12:70.56 Mb, 0.35, +1.05, 97% det), and MVK (chr12:109.63 Mb, 0.35, +1.40, 95% det) (Figure 9D, 9E).

The balanced oligodendrocyte candidates (Figure 9E, 9F, log2FC × strength) are ideal: 20 peaks clear log2FC ≥ 1 and strength ≥ 0.2, more than any other cell type in the atlas. The top ten include MVK (chr12:109.63 Mb, log2FC +1.40, 0.35 frags/cell, 95% det) — mevalonate kinase, the rate-limiting enzyme of the isoprenoid/cholesterol pathway that feeds myelin lipid synthesis; PREX1 (chr20:48.66 Mb, +1.04, 0.42 frags/cell, 95% det) — a Rac-GEF with known oligodendrocyte-lineage expression; SSNA1 (chr9:137.19 Mb, +1.28, 0.31, 97% det) — microtubule-regulating protein highly expressed in oligodendrocytes; PTPRB (chr12:70.56 Mb, +1.05, 0.35, 97% det); TECPR2 (chr14:102.42 Mb, +1.49, 0.25, 97% det); PRR18 (chr6:166.30 Mb, +1.10, 0.33, 97% det); DOHH (chr19:3.49 Mb, +1.10, 0.30, 100% det); TMC8 (chr17:78.11 Mb, +1.13, 0.27, 100% det); PCSK6 (chr15:101.52 Mb, +1.40, 0.21, 97% det); and MIR1265 (chr10:14.48 Mb, +1.60, 0.18, 97% det). TPRN, MAG, NUAK1, HAS1, and ATP5ME round out the top hits. Balanced candidates whose next-highest cell type is not OPC include CDC42EP2 (chr11:65.29 Mb, log2FC +1.89, 91% det, 0.44 frags/cell, next = TAM) and a distal SLC1A3 element (chr5:36.68 Mb, +2.15, 95% det, 0.22 frags/cell, next = TAM).

### Oligodendrocyte precursor cells (OPCs)

26 of 67 patients contribute ≥50 OPC cells. The pan-OPC distal regulatory landscape contains 318,959 peaks reliably open in ≥60% of OPC-evaluable patients, of which 42,261 are most open in OPCs and 5,248 are both reliable and selective (log2FC ≥ 1 vs the next cell type) (Figure 10A, 10B). No canonical OPC marker clears the 2× specificity threshold at the promoter peak. Only GPR17 (+0.06, 100% det), NEU4 (+0.01, 85% det), and PCDH15 (+0.37, 96% det) are OPC-primary at the promoter at all (Figure 10C). The second-highest cell type at OPC-selective distal peaks is excitatory neuron (41.6%), with GABA neurons at 26.8%, oligodendrocytes at 14.7%, and astrocytes at 11.8%.

**Figure 10.**
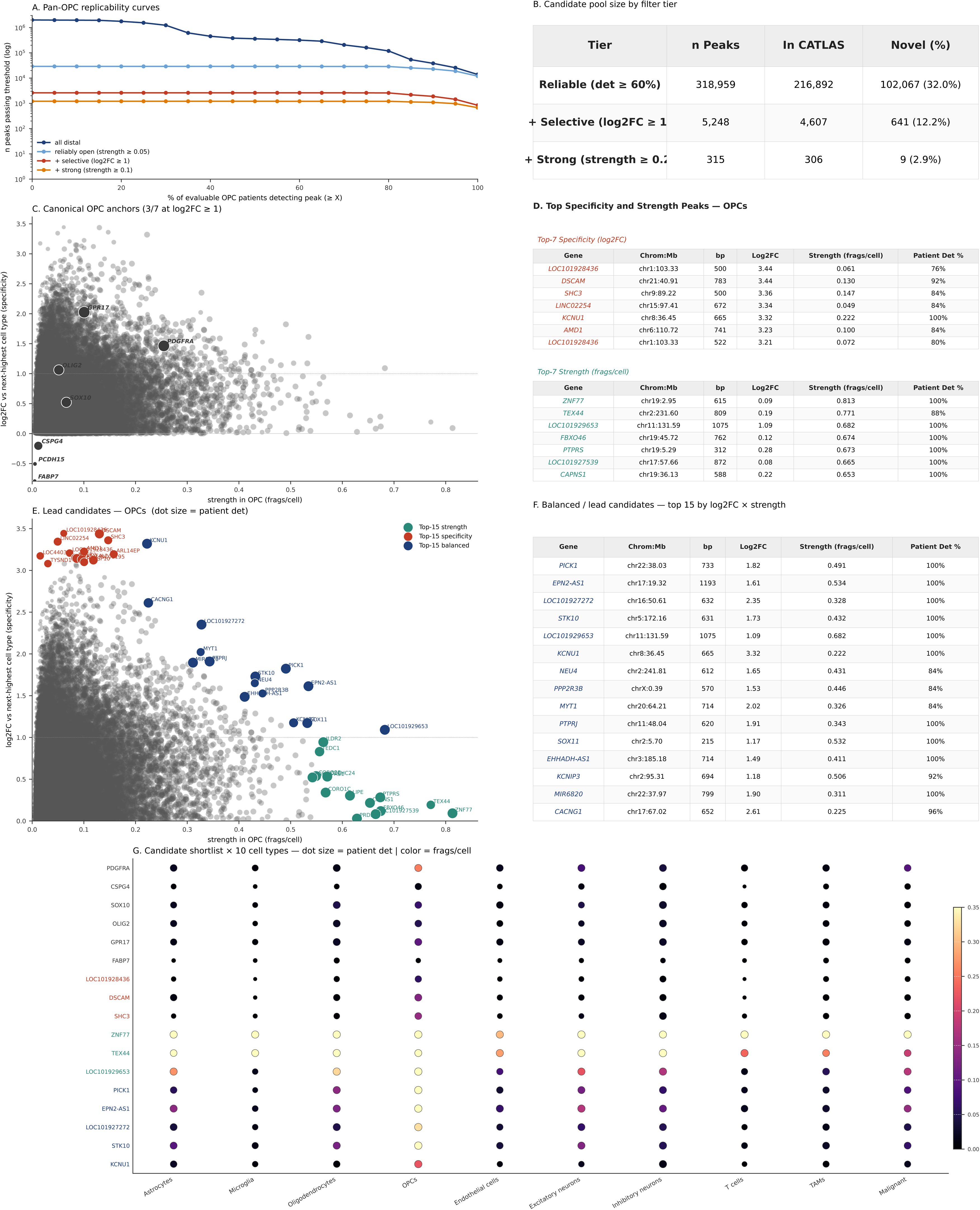
Oligodendrocyte precursor cell (OPC) candidate enhancers (26 patients with ≥50 OPC cells). (A) Replicability curves — number of distal peaks passing each patient-detection threshold, at four cumulative stringency tiers (all distal / reliably open at strength ≥ 0.05 / + selective at log2FC ≥ 1 / + strong at strength ≥ 0.1); log y-axis. (B) Candidate pool size table — three cumulative stringency tiers (reliable at det ≥ 60%; + selective at log2FC ≥ 1; + strong at strength ≥ 0.2), each with total n, number already in CATLAS, and number novel to CATLAS. (C) Canonical OPC lineage anchors in the strength × specificity space. X = strength in OPC (frags/cell, width-normalized); Y = log2FC vs next-highest primary cell type; dot size = patient detection rate. Charcoal dots = promoter peaks for the 7 canonical anchors (PDGFRA, CSPG4, SOX10, OLIG2, GPR17, FABP7, PCDH15); grey background = all OPC-primary distal peaks at det ≥ 60%. The panel headline reports how many anchors clear log2FC ≥ 1 (2/7 clear log2FC ≥ 1 for OPC). (D) Top Specificity and Strength Peaks tables, side by side — Top-7 highest-specificity (log2FC) candidates on the left, Top-7 highest-strength (frags/cell) candidates on the right. Columns: Gene, Chrom:Mb, bp, Log2FC, Strength (frags/cell), Patient Det %. (E) Lead candidates — combined specificity × strength view. X, Y, and dot size as in (C). Top-15 by strength (teal), top-15 by specificity (red), and top-15 by log2FC × strength composite (navy) are overlaid on the grey background of all OPC-primary distal peaks at det ≥ 60%. Horizontal dotted line = log2FC = 1 selectivity threshold. (F) Balanced / lead candidates — top 15 by log2FC × strength. Columns as in (D). (G) Candidate shortlist × 10 primary cell types. Rows = 6 canonical anchors + 3 top-specificity + 3 top-strength + 6 top-balanced candidates for OPC. Row label color encodes category (charcoal canonical / red specificity / teal strength / navy balanced). Dot size = patient detection rate; dot color = width-normalized strength.

Among the most specific OPC distal peaks are CHST15 (chr10:124.21 Mb, log2FC +2.98, 81% det, 0.08 frags/cell), LINC01195 (chr16:9.79 Mb, +2.76, 88% det, 0.12), LINC02254 (chr15:97.41 Mb, +2.65, 85% det, 0.05), LINC01648 (chr1:30.23 Mb, +2.63, 73% det, 0.06), and LINC00587 (chr9:102.50 Mb, +2.36, 92% det, 0.09) (Figure 10D, 10E). The highest-strength OPC peaks are headed by LOC101929653 (chr11:131.58 Mb, 0.68 frags/cell, log2FC +0.07, 100% det), ZCCHC24 (chr10:79.41 Mb, 0.57, +0.53, 100% det), KAZN-AS1 (chr1:14.26 Mb, 0.51, +0.79, 100% det), KCNIP3 (chr2:95.31 Mb, 0.51, +0.81, 92% det), and TOB2 (chr22:41.42 Mb, 0.50, +0.64, 100% det) (Figure 10D, 10E).

The balanced OPC candidates (Figure 10E, 10F, log2FC × strength) include BEGAIN (chr14:100.54 Mb, log2FC +1.50, 0.36 frags/cell, 100% det)—brain-enriched NMDA-receptor-associated protein; SLC15A4 (chr12:128.65 Mb, +2.07, 0.23, 100% det)—lysosomal peptide/histidine transporter; KIF19 (chr17:74.34 Mb, +1.06, 0.43, 100% det); HIP1R (chr12:122.85 Mb, +1.15, 0.36, 92% det); MMD2 (chr7:4.94 Mb, +1.04, 0.39, 100% det); LINC00844 (chr10:59.01 Mb, +1.68, 0.23, 100% det); ATCAY (chr19:3.89 Mb, +1.62, 0.23, 96% det); SGMS1 (chr10:50.51 Mb, +1.14, 0.30, 100% det); ZMIZ1-AS1 (chr10:79.06 Mb, +1.31, 0.26, 96% det); and LINC01195 (chr16:9.79 Mb, +2.76, 0.12, 88% det). GAS7, KIAA0040, LINC02517, COL13A1, and a second LINC00587 peak all have similarly high scores.

### T cells

T-cell regulatory elements could lend themselves to T-cell-modifying therapies, including in vivo CAR-T generation. 39 of 67 patients contribute ≥50 T cells to the analysis. T cells in GBM tissue are predominantly tumor-infiltrating lymphocytes (TILs), and the chromatin signatures below should be interpreted as TIL-derived30,31. The pan-T-cell distal regulatory landscape is relatively narrow, with 70,060 peaks reliably open in ≥60% of T-cell-evaluable patients, but with the highest selective fraction of any cell type: 6,086 distal peaks are most open in T cells and 2,809 are both reliable and selective (log2FC ≥ 1 vs the next cell type) (Figure 11A, 11B). Canonical marker promoters are the cleanest of any cell type in the atlas. A tier of pan-T TCR markers clears 2× at the promoter — CD3E (+3.10), CD3D (+2.48), CD3G (+2.48), LCK (+2.16), CD2 (+1.80), ZAP70 (+1.11) — alongside cytotoxic-effector markers PRF1 (+2.80), GZMA (+2.76), NKG7 (+2.29), GZMB (+1.75) and activation markers TIGIT (+1.34), CD4 (+1.23), CTLA4 (+1.03), EOMES (+1.02) (Figure 11C).

**Figure 11.**
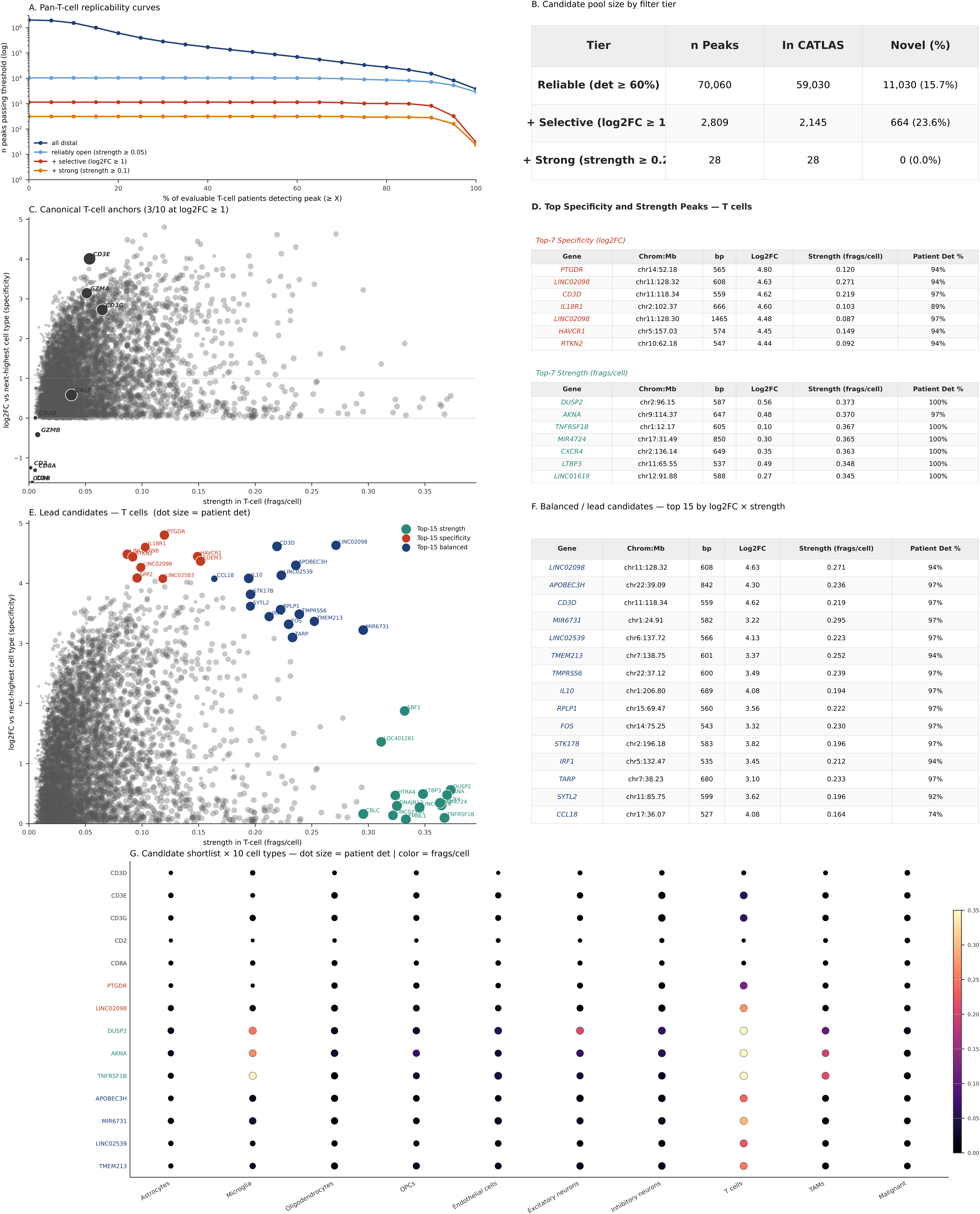
T cell candidate enhancers (3e patients with ≥50 T cells). (A) Replicability curves — number of distal peaks passing each patient-detection threshold, at four cumulative stringency tiers (all distal / reliably open at strength ≥ 0.05 / + selective at log2FC ≥ 1 / + strong at strength ≥ 0.1); log y-axis. (B) Candidate pool size table — three cumulative stringency tiers (reliable at det ≥ 60%; + selective at log2FC ≥ 1; + strong at strength ≥ 0.2), each with total n, number already in CATLAS, and number novel to CATLAS. (C) Canonical T cell lineage anchors in the strength × specificity space. X = strength in T cell (frags/cell, width-normalized); Y = log2FC vs next-highest primary cell type; dot size = patient detection rate. Charcoal dots = promoter peaks for the 11 canonical anchors (CD3D, CD3E, CD3G, CD2, TRAC, CD8A, CD8B, CD4, GZMA, GZMB, GNLY); grey background = all T cell-primary distal peaks at det ≥ 60%. The panel headline reports how many anchors clear log2FC ≥ 1 (7/11 clear log2FC ≥ 1 for T cell). (D) Top Specificity and Strength Peaks tables, side by side — Top-7 highest-specificity (log2FC) candidates on the left, Top-7 highest-strength (frags/cell) candidates on the right. Columns: Gene, Chrom:Mb, bp, Log2FC, Strength (frags/cell), Patient Det %. (E) Lead candidates — combined specificity × strength view. X, Y, and dot size as in (C). Top-15 by strength (teal), top-15 by specificity (red), and top-15 by log2FC × strength composite (navy) are overlaid on the grey background of all T cell-primary distal peaks at det ≥ 60%. Horizontal dotted line = log2FC = 1 selectivity threshold. (F) Balanced / lead candidates — top 15 by log2FC × strength. Columns as in (D). (G) Candidate shortlist × 10 primary cell types. Rows = 6 canonical anchors + 3 top-specificity + 3 top-strength + 6 top-balanced candidates for T cell. Row label color encodes category (charcoal canonical / red specificity / teal strength / navy balanced). Dot size = patient detection rate; dot color = width-normalized strength.

The strongest subtype-specific signals come from CD8+ cytotoxic markers (PRF1, GZMA, NKG7, GZMB) rather than CD4-helper markers (CD4 itself +1.23, IL7R +0.20, CCR7 +0.43). Treg-defining FOXP3 is essentially silent (18% patient detection, −3.08 log2FC), consistent with Tregs being a minor or absent fraction of GBM TILs. Among exhaustion markers, TIGIT (+1.34), EOMES (+1.02), BATF (+0.35) remain T-cell-primary at the promoter, while TOX (−1.52), LAG3 (−1.59), PDCD1 (−1.56), HAVCR2 (−0.91) are shared with neurons or myeloid cells — consistent with exhausted/dysfunctional TIL phenotypes in which these markers are also expressed in other GBM cell types.

Among the most specific T-cell distal peaks are CCL18 (chr17:36.07 Mb, log2FC +4.08, 74% det, 0.16 frags/cell), LINC02539 (chr6:137.72 Mb, +3.64, 97% det, 0.22), and SLC35G3 (chr17:35.21 Mb, +3.58, 92% det, 0.09) (Figure 11D, 11E). The highest-strength T-cell peaks are headed by DUSP2 (chr2:96.15 Mb, 0.37 frags/cell, log2FC +0.35, 100% det), AKNA (chr9:114.37 Mb, 0.37, +0.48, 97% det), and TNFRSF1B (chr1:12.17 Mb, 0.37, +0.10, 100% det) (Figure 11D, 11E).

The balanced T-cell candidates (Figure 11E, 11F, log2FC × strength) are driven by exceptionally clean pan-T cell signal. The top ten include LINC02098 (chr11:128.32 Mb, log2FC +3.23, 0.27 frags/cell, 95% det); LINC02539 (chr6:137.72 Mb, +3.64, 0.22, 97% det); a CD3D distal element (chr11:118.34 Mb, +3.49, 0.22, 97% det); MIR6731 (chr1:24.91 Mb, +2.53, 0.30, 97% det); CCL18 (chr17:36.07 Mb, +4.08, 0.16, 74% det); SYTL2 (chr11:85.75 Mb, +3.40, 0.20, 92% det); TMPRSS6 (chr22:37.12 Mb, +2.77, 0.24, 97% det); and APOBEC3H (chr22:39.09 Mb, +2.80, 0.24, 97% det).

### Excitatory neurons

44 of 67 patients contribute ≥50 excitatory neurons (the non-GABAergic cortical neuron pool; in cortex this is >90% glutamatergic, which we have simplified here to excitatory neurons). This count exceeds the patient count for most cell types because a substantial fraction of contributing samples came from marginal/peritumoral tissue with low tumor cellularity and correspondingly higher neuron content. One patient’s excitatory-neuron data (dejong_2026 patient AT9, n = 265 cells) was excluded from the neuron set for being a marked multi-chromosomal outlier, contributing an unreasonable 46−280× the next-highest per-cell fragment density at every suspect locus (chr1, chr7, chr12, chr16) while canonical-marker peaks showed normal signal. The pattern is consistent with a small cluster of CNV-heavy malignant cells mis-labeled as “neuron” during annotation; no other pseudobulk in any cell type showed comparable contamination. After exclusion, the pan-excitatory-neuron distal landscape contains 240,119 peaks reliably open in ≥60% of the remaining 44 neuron-evaluable patients, of which 107,385 are most open in excitatory neurons and 33,585 are both reliable and selective (log2FC ≥ 1 vs the next cell type). Top-strength and balanced candidates sit in the 0.2−0.9 frags/cell range (max 0.92).

Among the most specific excitatory-neuron distal peaks are GABRG3-AS1 (chr15:27.46 Mb, log2FC +5.20, 64% det, 0.22 frags/cell), PASK (chr2:241.16 Mb, +5.19, 66% det, 0.30), LINC02670 (chr10:9.92 Mb, +4.88, 64% det, 0.21) (Figure 12E1). Patient detection across this top-specificity tail sits in the 60−70% range, close to the atlas’s 60% reliability gate. The highest-strength excitatory-neuron peaks are headed by PER1 (chr17:8.16 Mb, 0.92 frags/cell, log2FC +0.17, 95% det), DBNDD1 (chr16:90.02 Mb, 0.87, +0.67, 93% det), and CLEC11A (chr19:50.73 Mb, 0.87, +0.82, 70% det).

**Figure 12.**
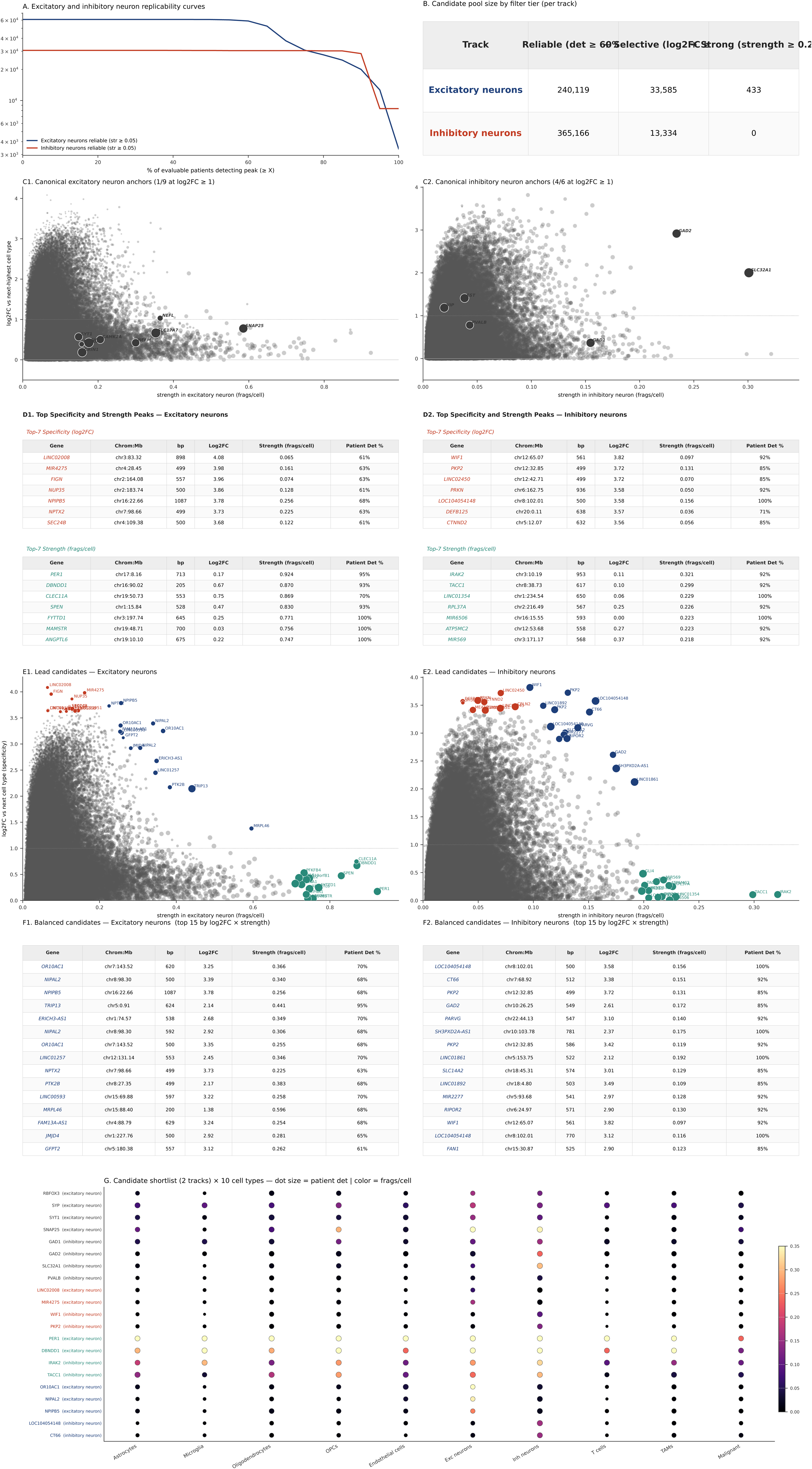
Excitatory and inhibitory neuron candidate enhancers. Two tracks: excitatory neurons (44 patients with ≥50 cells) and inhibitory neurons (14 patients with ≥50 cells). Dejong 2026 patient AT9 was excluded from the excitatory track as a CNV-heavy outlier (Methods). (A) Replicability curves — number of distal peaks reliably open (strength ≥ 0.05) in each track, as a function of patient-detection threshold. Two curves overlaid (navy excitatory, red inhibitory); log y-axis. (B) Candidate pool size table — two tracks × three cumulative stringency tiers (reliable at det ≥ 60%; + selective at log2FC ≥ 1; + strong at strength ≥ 0.2), each with total n, number already in CATLAS, and number novel to CATLAS. (C1–C2) Canonical lineage anchors in the strength × specificity space, one scatter per track. Excitatory anchors (C1): RBFOX3, SYP, SYT1, SNAP25, NEFL, NEFM, SLC17A7, GRIN1, CAMK2A. Inhibitory anchors (C2): GAD1, GAD2, SLC32A1, PVALB, SST, VIP. Charcoal = canonical anchors; grey background = all primary-track distal peaks at det ≥ 60%; dot size = patient detection rate. Panel headlines report how many anchors clear log2FC ≥ 1 (1/9 excitatory, 4/6 inhibitory). (D1–D2) Per-track Top Specificity and Strength Peaks — Top-7 highest-specificity (log2FC) above, Top-7 highest-strength (frags/cell) below, each as a mini-table with columns Gene, Chrom:Mb, bp, Log2FC, Strength (frags/cell), Patient Det %. (E1–E2) Per-track lead candidates — combined strength × log2FC overlay. Top-15 by strength (teal), specificity (red), and balanced composite (navy) over the grey per-track background. Horizontal dotted line = log2FC = 1 selectivity threshold. (F1–F2) Per-track balanced top-15 candidate tables — gene, Chrom:Mb, bp, log2FC, strength, patient detection. (G) Combined candidate shortlist dotplot — 4 canonical anchors + 2 top-specificity + 2 top-strength + 3 top-balanced per track × 10 primary cell types. Row labels carry the originating track in parentheses (excitatory or inhibitory); label color encodes category (charcoal canonical / red specificity / teal strength / navy balanced). Dot size = patient detection rate; color = width-normalized strength.

The balanced excitatory-neuron candidates (Figure 12F1, 12G, log2FC × strength) include OR10AC1 (chr7:143.52 Mb, log2FC +3.25, 0.37 frags/cell, 70% det), NIPAL2 (chr8:98.30 Mb, +3.39, 0.34, 68% det), NPIPB5 (chr16:22.66 Mb, +3.78, 0.26, 68% det), TRIP13 (chr5:0.91 Mb, +2.14, 0.44, 95% det), ERICH3-AS1 (chr1:74.57 Mb, +2.68, 0.35, 70% det), LINC01257 (chr12:131.14 Mb, +2.45, 0.35, 70% det), NPTX2 (chr7:98.66 Mb, +3.73, 0.23, 64% det) — neuronal pentraxin 2, an immediate-early gene classically expressed in cortical excitatory neurons — and PTK2B (chr8:27.35 Mb, +2.17, 0.38, 68% det). Patient detection for most balanced candidates sits in the 64−70% range (only TRIP13 reaches ≥ 90%), reflecting the heterogeneity of neuron content across contributing samples.

### Inhibitory neurons

14 of 67 patients contribute ≥50 inhibitory neurons to the analysis. The open chromatin signal is markedly different from excitatory neurons: 365,166 peaks are reliably open in ≥60% of inhibitory-neuron-evaluable patients, of which 91,299 are most open in inhibitory neurons and 13,334 are both reliable and selective (log2FC ≥ 1 vs the next cell type). Canonical inhibitory markers cluster at modest specificity: GAD2 (+2.81, 93% det, 0.23 frags/cell) and TAC1 (+2.03, 100% det) clear the 2× threshold at the promoter; SLC32A1/VGAT (+0.92), VIP (+0.93), SST (+0.90), PVALB (+0.49), DLX6 (+0.14), DLX5 (+0.27) remain inhibitory-primary at the promoter at low fold-change. GAD1 (−0.94), RELN (−0.12), LHX6 (−0.03), NPY (−0.67), CALB1 (−0.87), and CALB2 (−0.38) score fractionally higher in excitatory neurons at the promoter peak (Figure 12C2).

Among the most specific inhibitory-neuron distal peaks are SH3BGRL (chrX:81.78 Mb, log2FC +3.72, 64% det, 0.04 frags/cell), LOC104054148 (chr8:102.01 Mb, +3.48, 100% det, 0.16), and LOC101927829 (chr3:20.56 Mb, +3.38, 86% det, 0.07) (Figure 12E2). The highest-strength candidates are modest (0.21−0.25 frags/cell) and the top-strength tail does not include any 2×-specific peaks.

The balanced inhibitory-neuron candidates (Figure 12F2, 12G) include LOC104054148 (chr8:102.01 Mb, log2FC +3.48, 0.16 frags/cell, 100% det); CT66 (chr7:68.92 Mb, +3.03, 0.15, 93% det); a GAD2 distal element (chr10:26.25 Mb, +2.64, 0.17, 86% det) — adjacent to the canonical GABA-synthesis marker; LINC01861 (chr5:153.75 Mb, +2.13, 0.19, 100% det); SH3PXD2A-AS1 (chr10:103.78 Mb, +2.23, 0.18, 100% det); and CDH9 (chr5:26.94 Mb, +2.15, 0.16, 86% det).

### Therapy-focused features

Across all cell types, we examined how well the top 1000 peaks replicated across patients. After confining analysis to peaks present in at least 60% of patients, oligodendrocyte, T cell, OPC, inhibitory neuron, and microglia top-1000 candidates typically reach ≥85% of evaluable patients (medians 84−93%); astrocyte and endothelial candidates reach a lower fraction (medians 70−81%); malignant and excitatory neuron candidates cluster closer to the 60% reliability floor (Figure 13A). Cell types with fewer evaluable patients (inhibitory neuron (n=14), microglia (n=19), and endothelial (n=20)) carry greater uncertainty in these distributions, and the small-sample medians should be treated as provisional.

**Figure 13.**
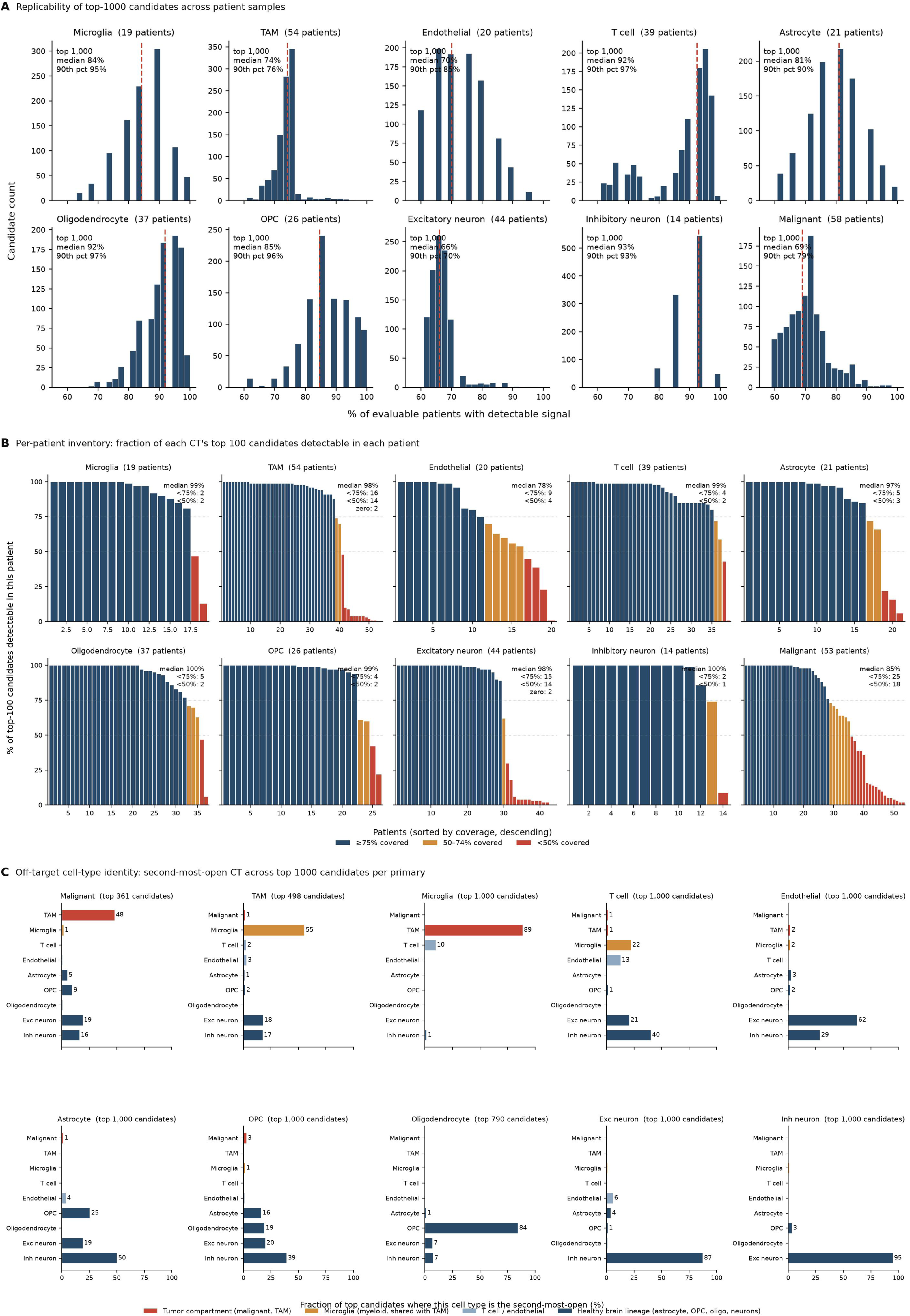
Candidate replicability and off-target profile across patient samples. Summary views of the atlas’s candidate pool framed around considerations for cell-type-targeted AAV therapy. (A) Patient-level replicability of the top-1000 candidates by primary_score for each primary cell type (after filtering to det ≥ 60%). Histograms show, across top-1000 candidates per cell type, the fraction of evaluable patients with detectable signal in each candidate; red dashed line marks the median. Inset values: top-N (or "all qualifying" where fewer than 1000 peaks pass the reliability filter), median patient coverage, and 90th-percentile patient coverage. Panel titles show the number of evaluable patients for that cell type. (B) Per-patient candidate inventory: for each patient sample of each primary cell type, the fraction of that cell type’s top-100 candidates that are detectable (≥ 1 fragment) in that patient. Bars sorted by coverage per cell type, descending. Color tiers: navy ≥ 75%, amber 50–74%, red < 50%. Dashed horizontal guides at 75% and 50%. Panel insets report median patient coverage and counts of patients below each threshold. (C) Off-target identity across the top-1000 candidates for each primary cell type: horizontal bars show, for each candidate, the second-most-open cell type (the cell type with the highest non-primary strength). Bar colors denote the broad tissue compartment of the off-target: red = tumor compartment (malignant, TAM); amber = microglia (myeloid lineage, shared with TAM); light blue = T-cell or endothelial; navy = healthy brain lineage (astrocyte, OPC, oligodendrocyte, excitatory or inhibitory neuron).

Patient heterogeneity is a point faced by every clinical trial, and a point frequently raised when working with human patient samples. We therefore took each cell type’s top 100 peaks and asked whether it could be expected to work in a given patient. In other words, how well do these peaks apply to each individual patient? For six of ten cell types (oligodendrocyte, inhibitory neuron, T cell, microglia, OPC, astrocyte), the median patient sample contains ≥97% of the top-100 peaks (Figure 13B). Such a designed candidate panel would reach essentially every patient in the atlas. For TAM and excitatory neuron, median coverage is similarly high (98%) but a tail of patients (14 of 54 and 14 of 44, respectively) detects fewer than half of the top-100, and two patients in each show no detectable candidate at this pool size. Malignant and endothelial show substantially more per-patient variability (medians 85% and 78%; one-third or more of patients fall below 75% coverage).

Two factors contribute to these low-coverage patients: inter-patient biological heterogeneity, particularly for malignant chromatin configurations, and per-patient fragment sequencing depth within a cell-type-specific profile, which caps the number of rare peaks any patient can recover regardless of candidate choice.

Finally, for each cell type’s top 1000 peaks, we tabulated the highest off-target cell types (Figure 13C). TAM and microglia top candidates are predominantly second-ranked in each other (tolerable for myeloid-directed therapy), while oligodendrocyte top candidates have OPCs as the dominant second-rank (a potential safety concern depending on payload).

## Discussion

Cell-type-specific enhancers offer the ability to manipulate individual cell types selectively. Significant effort has been expended developing such tools for mouse and other mammalian brains^3-13^. Since two-thirds of such sequences in humans are not conserved, it is vital that we use data from human brain cell types to identify such potential tools for human therapies^14^. Moreover, many cell types in neurological pathologies will not retain the exact chromatin accessibility patterns found in normal human brain cell types; using human brain tissue specifically from these pathologies is therefore imperative.

In order to begin addressing this, we present the first atlas harmonizing all publicly available human IDH-wildtype glioblastoma snATAC-seq data, assembling 67 patients across ten studies^15-24^. This atlas establishes that the chromatin landscape of human glioblastoma encodes a reproducible, distal enhancer repertoire for every major cell type in the tumor and its microenvironment, and that many cell-type-selective regulatory elements in tumor-involved compartments are absent from normal-brain references.

The strongest selective signal for nearly every cell type is at distal elements, consistent with prior work^25,26^. We therefore restricted the candidate set to peaks ≥2 kb from any annotated transcriptional start site. Excitatory and inhibitory neurons yield the largest pools of reliable, selective distal peaks (tens of thousands each); microglia and OPCs yield several thousand; astrocytes, T cells, and endothelial cells yield hundreds to low thousands within smaller evaluable patient sets. The narrowest pools belong to oligodendrocytes, TAM, and malignant cells, a few hundred candidates each, despite these cell types contributing the largest numbers of evaluable patients in the atlas. The constraint for these three is chromatin overlap: oligodendrocyte chromatin shares extensively with other glial lineages, TAM with microglia (shared myeloid lineage), and malignant with multiple non-malignant compartments owing to transcriptional-state heterogeneity. T cell and oligodendrocyte top candidates show the highest patient-level convergence in the atlas (median 92% of evaluable patients).

CATLAS, the current reference adult human brain snATAC-seq resource, was built with post-mortem adult brain samples. The glioma-involved microenvironment (TAM, reactive microglia, tumor vasculature, malignant cells) is absent from these, so a disease atlas is a prerequisite for targeting such tumor-involved compartments. ∼78% of TAM peaks and ∼84% of endothelial cell peaks are novel to CATLAS, highlighting the regulatory biology of cell types closely involved with glioblastoma, though notably malignant and microglia selective peaks almost completely overlap CATLAS peaks (0.3% and 0% novel, respectively). A synthetic super-enhancer construct assembled from fragments screened in cultured patient-derived GSC lines produced curative outcomes in preclinical GBM models and is advancing to clinical trial^28^. The cross-patient, cell-type-resolved atlas presented here, derived from primary human GBM tissue, is the foundation for programs that extend beyond the GSC compartment to all involved cell types. Other diseases with important pathogenic cell types would similarly benefit from such patient-derived atlases.

This atlas carries several limitations. An atlas like this is limited to publicly existing data, which remains sparse and in need of further well-defined and deeply sequenced samples. Some thousands of surgeries every year resect glioblastoma with tissue in excess of what is needed for pathological diagnosis, but only 67 patient samples have been processed with publicly available single-nucleus ATAC sequencing. Many of these do not offer public access to fragment-level files. Copy-number variation inflates apparent chromatin accessibility on gained chromosomes and deflates it on lost ones; 61% of the naive pan-malignant top-100 candidates show copy-number artifacts, predominantly at the EGFR locus. We flag rather than correct these because a principled per-patient adjustment would require fragment-level data that six of ten cohorts did not make publicly accessible; users with copy-number-sensitive applications may wish to restrict analysis to the four fragment-mode cohorts, or apply their own correction. Malignant subtype resolution is incomplete: Neftel class assignment is available only in the largest cohort, and the pan-malignant pool necessarily averages across transcriptional states whose chromatin may diverge.

Finally, this atlas can provide only candidate enhancer sequences—predictions of which sequences will drive sufficient and sufficiently selective expression is difficult, though in some cases up to a third or more of candidates are useful^4,5,7,10,11^. Animal models are known to poorly model the human disease, and mouse genomes lack a large proportion of human enhancers. Next to studies in actual human patients, the next best functional validation approach for these types of sequences is in *ex vivo* glioblastoma tissue. This is the natural next step for this work.

## Methods

We intend this atlas to be a continually updated work in progress. We will add future studies as they become available, and look forward to incorporating any suggestions and modifications that can make it more useful to the general community.

### Cohort inclusion criteria

We assembled every publicly accessible human adult IDH-wildtype glioblastoma snATAC-seq or 10x Multiome dataset for which raw fragments or processed accessibility matrices were available at the time of build freeze. We excluded patient-derived cell lines, patient-derived organoids and xenografts, glioma stem cell cultures, flow-sorted fractions, pediatric high-grade glioma, diffuse midline glioma, and tumors with IDH1/2 mutations as defined by the WHO 2021 GBM criteria. Primary and recurrent samples were both retained; recurrence status is recorded in per-sample metadata. Cohorts for which raw fragments were unavailable but author-processed matrices were open were included (see "Per-cohort ingest"). Controlled-access deposits, data behind unresolved data-use agreements, and deposits on sequencing platforms for which no in-house ingest adapter was available were excluded from the current build and are listed with their accession and reason in the cohort-summary table.

Ten cohorts cleared inclusion: de Jong et al. 2026 Nature (E-MTAB-17183), Sojka et al. 2025 Nature Cell Biology (GSE276177), Sundaram et al. 2024 Science (GDC open-access pan-cancer scATAC blob, UUID a7765b5f-a606-4657-94ed-4c2a85bf3f39), Lin et al. 2026 Nature Neuroscience (HRA004942, CNCB-GSA), Wang X et al. 2024 Science Advances (CNP0003766, CNGB), Wang L et al. 2019 Cancer Discovery (GSE138794), Guilhamon et al. 2021 eLife (GSE139136), Raviram et al. 2023 PNAS (GSE165037), Pai et al. 2025 Cancer Research (GSE286415), and Terekhanova et al. 2023 Nature (GSE240822). Full per-cohort provenance is in the cohort-attribution table deposited with the atlas. After cell-level quality control, the atlas contains 67 patients with at least 50 cells per labelled cell-type group across 10 cohorts and 1,650,376 nuclei.

For Pai 2025, the sample-naming divergence between GEO fragment filenames (e.g. GBM{N}M, GBM3A/B, GBM5A/B) and the author metadata table (e.g. GBM{N}I, GBM3C/I, GBM5C/I) was reconciled by a per-sample barcode-overlap check against 50,000 lines from each fragments file; the IDH-mutant GBM2 patient was excluded per the author-supplied IDH status table; the ambiguous OL-MG oligo-microglia cross-lineage class was dropped; and the Myeloid label was mapped to TAM because the deposit did not distinguish tumor-associated macrophages from microglia, and the tumor-core and margin contexts default to TAM. For de Jong 2026, which labelled all brain-resident and infiltrating myeloid cells into eleven TAM subtypes without a separate microglia class, the authors’ Resident-TAMs subtype (8,236 cells across 12 patients) was transferred to the microglia pool; the remaining TAM subtypes stayed in the TAM pool. The transfer was validated against a six-marker ATAC signature at ±10 kb windows around canonical microglia markers (CX3CR1, TMEM119, P2RY12, HEXB, SALL1, CSF1R) versus macrophage markers (CD163, MRC1, APOE, TYROBP, F13A1, LYVE1); Resident-TAMs cells scored 66% microglia-leaning per cell versus 35% in Resident BAM TAMs and 4−7% in non-myeloid controls. A pan-myeloid track combining microglia + TAM + BAM is retained as an additional per-cell-type aggregate for cross-cohort analyses in which the microglia/TAM distinction is weak.

### Per-cohort ingest

Every cohort was loaded into a common per-cell data object with fixed metadata (cohort, sample, patient, modality, IDH status, primary-vs-recurrent status, anatomical region). Three ingest paths were used, depending on what the source deposit contained:

#### Fragment-mode

Cohorts that released raw per-cell Tn5 fragment files on GEO or GDC were loaded directly from fragments, and per-cell quality metrics (TSS enrichment, fragment-size distribution, fragments per cell) were computed de novo against hg38. This path is the most permissive downstream because every per-cell measurement is re-constructible. It applies to the Sojka 2025, Sundaram 2024, Pai 2025, and Terekhanova 2023 cohorts.

#### Matrix-mode

Cohorts that released only author-processed peak × cell count matrices were lifted onto the atlas coordinate frame at the quantification step. Per-cell fragments are not reconstructible and TSS enrichment is unavailable. This path applies to de Jong 2026, Lin 2026, Wang X 2024, Wang L 2019, Guilhamon 2021, and Raviram 2023. Cohorts whose deposited artefact was an ArchR, Signac, or SnapATAC R object (Raviram 2023 as sci-ATAC output) were converted to a Python-readable form before entering the pipeline.

### Quality control

Two per-cell gates were applied in series for fragment-mode cohorts: TSS enrichment of at least 6 (snapatac2 default) and fragments per cell between 1,000 and 100,000. For matrix-mode cohorts, where TSS enrichment is not available, we applied the fragment-count floor on non-zero peak counts per cell at the same 1,000 cutoff. At the donor level, we removed any donor whose mean peaks per cell fell below 500, because at that depth peak calling and specificity scoring become dominated by sampling noise. Donor-level and per-cell QC pass rates are reported in the cohort-summary table. No central doublet caller was run in this build. We relied on each contributing cohort’s upstream doublet removal, together with the TSS-enrichment and fragment-count gates, and two cohort-specific removals of author-flagged cross-lineage ambiguity classes (Pai 2025’s OL-MG class, and any cell whose author annotation was itself a doublet label).

### Cell type labeling

#### Malignant-cell calling

Tumor cells were called per cohort using inferCNVpy on gene activity, computed from Tn5 insertion density across hg38 gene bodies for fragment-mode cohorts, or taken directly from the paired RNA matrix for the de Jong 2026 multiome. A cell was called malignant if its CNV score exceeded μ + 2σ of its own sample’s reference distribution. The CNV panel was the standard IDH-WT GBM arm-level fingerprint: +chr7 (EGFR), −chr10 (PTEN), −chr9p (CDKN2A/B), +chr20, +chr19. The sliding 100-gene window is deliberately coarse so the score reflects whole-arm events; focal amplifications are flagged separately at the scoring layer. We also computed a chr7+/chr10− accessibility ratio — (mean chr7 + ε) / (mean chr10 + ε) ≥ 2.0 — which agreed with the inferCNV call on 86−93% of cells in fragment-mode cohorts. For matrix-mode cohorts without gene-level resolution, this ratio was used directly as the malignant flag.

#### Author passthrough

Cohorts whose deposits carried a usable per-cell annotation (de Jong 2026 TAM/BAM subtypes; Pai 2025 CellType; Wang L 2019 author labels) used author labels as the ground truth for cell types after the cohort-specific remapping described above. A standardized mapping translated each author vocabulary into the atlas ontology, which has 10 primary cell types (astrocyte, microglia, oligodendrocyte, OPC, endothelial, neuron, GABA_neuron, T_cell, TAM, malignant pooled) and 4 malignant Neftel subtypes (AC-like, MES-like, NPC-like, OPC-like). Each atlas cell type is mapped to a Cell Ontology (CL) term for cross-reference against CELLxGENE and Human Cell Atlas resources: astrocyte CL:0000127; microglia CL:0000129; oligodendrocyte CL:0000128; OPC CL:0002453; endothelial cell CL:0000115; excitatory (glutamatergic) neuron CL:0000679; inhibitory (GABAergic) neuron CL:0000617; T cell CL:0000084; tumor-associated macrophage CL:0000864 (parent macrophage); malignant cell CL:0001064 (malignant cell; Neftel subtypes retain the parent term with an atlas-local subtype annotation).

#### scRNA-bridged label transfer

For cohorts with gene-level signal but no workable author annotation, cell type labels were transferred from GBmap (a disease-matched human GBM scRNA reference of 109 patients and 330,000 cells^29^) using scGLUE for ATACXRNA cross-modal mapping where it converged, with a CellTypist fallback on gene-activity scores where scGLUE did not converge. Neuronal subtypes were transferred from CATLAS (107 adult human brain cell types^14^) by nearest-neighbor in the shared peak-space.

#### Marker-peak scoring

For cohorts without gene-level signal, each union peak was mapped to any gene whose body ±5 kb overlapped the peak midpoint. A per-cell signature was computed per cell type as the mean z-score of accessibility across that cell type’s marker set, and a cell was assigned to its best-scoring type when the per-cell maximum z was ≥1.0 and the margin over the runner-up was ≥0.3; otherwise the cell was labelled unassigned. The canonical marker sets are in Supplementary Table 2.

#### Re-annotation of coarse author labels

For Guilhamon 2021 and Lin 2026, which deposited only coarse malignant versus TAM/unassigned annotations, the marker-peak scoring step was applied on top of the author label to recover astrocyte, oligodendrocyte, OPC, neuron, endothelial, and T-cell calls where the signature cleared the two-criterion rule. Existing granular labels were preserved; only unassigned-like labels were replaced. Any cell called malignant by CNV was routed to the Neftel subtype scorer regardless of what other label systems returned; it never entered the non-malignant path. This prevents CNV-defined tumor cells from being mislabeled when their accessibility is incidentally open at a non-malignant marker promoter. For non-malignant cells, author passthrough took precedence over GBmap, GBmap over CATLAS, and CATLAS over marker-peak scoring, with the exception that CATLAS neuronal subtype calls overrode a generic GBmap neuron call. Cells called malignant were scored on the four Neftel 2019 marker modules (AC-like, MES-like, NPC-like, OPC-like) using marker-set scoring on gene-activity space and assigned to the highest-scoring module. For cohorts in peak-space with no addressable Neftel marker genes, malignant cells were tagged malignant_unresolved and entered the pan-malignant pool rather than a subtype pool.

### Union peak set construction

Per-cell-type pseudobulk Tn5 insertion sets were generated for every cohort × patient × cell-type group by subsetting per-cell fragments (fragment-mode cohorts) or by inheriting peak intervals from the source matrix. MACS3 was run on each fragment-mode pseudobulk in paired-end mode with the ATAC-seq-standard shift/extend parameters and a q-value floor of 0.05, producing per-cell-type narrow-peak sets per cohort. The atlas union peak set was constructed by combining CATLAS 544K adult-brain peak atlas with the novel MACS3 peaks called on every fragment-mode cohort, merged across sources without transitive chaining across cohorts. The resulting union contains 2,085,175 intervals (mean width 594 bp, maximum 6.7 kb). Per-peak contributing-source provenance is retained at single-interval resolution, so any downstream user can recover which cohorts and which caller contributed to any union peak.

#### Pseudobulk construction

Fragment-mode cohorts were re-quantified against the union peak set using snapatac2’s peak-matrix routine on each cohort’s own fragments. Matrix-mode cohorts were lifted onto the union set by counting overlap-weighted fragments where a source peak intersected a union peak; for sources whose peak set was coarser than a union peak, the source signal was apportioned across overlapping union peaks by overlap length. Pseudobulk counts were then formed for every (cohort, patient, cell type) group with at least 50 cells after QC. We kept per-peak CPM (counts per million, normalized by the pseudobulk’s total count across union peaks) and per-cell strength, defined as the mean Tn5 fragment count per cell overlapping the peak, width-normalized to a 500 bp reference peak (multiplied by 500/peak_width_bp) so that strength values are comparable across peaks of different widths. Both CPM and per-cell strength were kept because they answer different questions: CPM is a compositional readout of where a cell type’s sequencing library lands; per-cell strength is a density readout of how often, per cell, Tn5 inserts at this locus. Downstream, per-cell strength is the primary quantity for AAV enhancer selection, because an AAV capsid at a locus experiences a per-cell density of accessibility, not a compositional fraction of the cell type’s total library.

### Enhancer scoring

#### Specificity

We define specificity as the log2 ratio of per-cell strength in the peak’s top cell type to its next-highest cell type across the full atlas taxonomy, with a small constant (1e-6) added to numerator and denominator to prevent division by zero. The next-highest cell type (rather than pooled background, the ArchR default) we think is the better denominator for potential AAV therapeutic design because the major problem would be a secondary off-target hit that a grouped median/average would mask.

#### Strength

The average per-cell fragment count is calculated over every evaluable patient, not only patients in which the peak is detected, to avoid strength being computed on a patient subset defined by a detection threshold.

#### Within-atlas replicability

For each peak × cell type, the number of evaluable patients in which the peak was detected, divided by the number of evaluable patients, where detection requires at least 1% of that patient’s cells of the target type to carry ≥1 Tn5 fragment at the peak. A hard floor of 60% within-atlas patient detection was applied before the top-N selection used in the shipped per-cell-type candidate tables.

#### Cross-cohort replication

The number of the ten cohorts in which the peak cleared a per-cohort minimum detection threshold (≥5% of cells in that cohort with ≥1 fragment at the peak). Patient reproducibility and cross-cohort study replication are reported independently.

#### Primary ranking

The default ranking column is the mean of the distal (>2kb from TSS) percentile ranks of the specificity and strength components. Rank-percentile formulation is nonparametric and makes no assumption about the scale or distribution of either component; it penalizes near-zero specificity directly, because a near-zero log2FC lands in a near-zero percentile.

#### Taxonomy separation

The primary cell-type tabs compute specificity across the 10 primary cell types (pooled malignant counted as one lineage; Neftel subtypes excluded from the denominator). The malignant-subtype tabs compute specificity across 13 cell types (9 non-malignant plus 4 Neftel subtypes). The pan-myeloid grouping similarly recomputes a pooled myeloid group against all other cell types.

#### Pan-malignant aggregate and amplicon-adjacent flagging

A separate pan-malignant per-peak × per-patient × per-cohort aggregate was built to group all tumor cells together, bypassing Neftel sub-typing. A cell entered the pan-malignant pool if it was called malignant by CNV OR carried the malignant_unresolved label. The CNV-flag arm of the OR is necessary because marker-peak scoring in matrix-mode cohorts can label CNV-malignant cells as TAM, microglia, OPC, or neuron whenever the tumor cell’s chromatin incidentally opens at a cell-type marker TSS; the de Jong cohort alone contributes 438,994 CNV-called malignant cells. Per-patient aggregates require at least 10 malignant cells per patient and apply a 10% within-patient accessibility threshold for consistency scoring. Peaks within ±1 Mb of four canonical GBM amplicon or deletion loci are flagged in every candidate table with an AMP designation and the responsible locus: EGFR (chr7:54.5−56 Mb), MDM2 (chr12:68−70 Mb), MYC (chr8:127−129 Mb), and CDKN2A (chr9:21−22.5 Mb). The flag is informational: rows remain in all tables and rankings.

### Programmatic access and AI usability

#### Column schema

Every per-cell-type candidate table and the full per-peak master-score parquet use a fixed column schema so that external pipelines and LLM agents can query without inspection. Each row is one union peak. Coordinate and annotation columns: peak_id (primary key), chrom / start / end / peak_width_bp / is_distal (hg38), nearest_gene and dist_to_tss_signed (RefSeq, nearest-gene body ±5 kb). Top-level fields: top_celltype_primary names the strength-argmax cell type (over the 10 primaries), and log2FC_vs_primary gives that peak’s log2 fold-change against the next-highest cell type. Per-track log2 fold-change is also persisted for the pan-myeloid aggregate (log2FC_myeloid) and each of the four Neftel malignant subtypes (log2FC_malignant_AC, log2FC_malignant_MES, log2FC_malignant_NPC, log2FC_malignant_OPC). For each of the 10 primary cell types, the pan-myeloid aggregate, and the four Neftel subtypes (15 cell types in total, suffix _{C} in the column name), the following metric columns are present: strength_frags_per_cell_wn_{C} (per-cell width-normalized Tn5 fragments), det_rate_{C} (fraction of evaluable patients detecting the peak in that cell type), n_eval_{C} (patients contributing ≥50 cells of that cell type), n_det_{C} (patients in whom the peak was detected), n_cohorts_{C} (distinct cohorts in which the peak replicated at ≥5% within-cohort detection), cpm_{C} (compositional counts-per-million), primary_score_{C} (mean of strength and specificity rank percentiles, the default ranking column), and strength_pct_{C} / spec_pct_{C} (per-cell-type rank percentiles). CATLAS overlap and amplicon-adjacency flags are not persisted in the parquet; they are computed on demand from the standalone reference files supplied alongside the atlas (reference/catlas_peaks.bed from Li YE 2023 Science, and reference/gbm_amp_loci.bed for the four flagged amplicon/deletion windows described in the amplicon-flagging paragraph above).

#### Quick-start query

The pseudobulk master-score parquet is designed for direct pandas or polars query without additional joins. For example, the top 50 microglia-selective distal candidates with cross-cohort replication in ≥5 cohorts are extracted in three lines: *import pandas as pd df = pd.read_parquet(’*https://mcginnislab-gbm-atlas.s3.amazonaws.com/reports/atlas_v4/v4_master_scores.parquet*’) df.query(’top_celltype_primary == "microglia" and det_rate_microglia >= 0.60 and log2FC_vs_primary >= 1 and n_cohorts_microglia >= 5’).nlargest(50, ‘primary_score_microglia’)*

An R-tidyverse equivalent using arrow::read_parquet is one line longer and is included in the repository’s examples/ folder along with a Python notebook that reproduces every per-cell-type candidate table from scratch.

### Reproducibility and data/code availability

Data are permanently archived at Zenodo (concept DOI: 10.5281/zenodo.23252323; this analysis used version v2026-11-15, DOI 10.5281/zenodo.24270756) and hosted on the AWS Registry of Open Data (https://registry.opendata.aws/gbm-chromatin-accessibility-atlas-mcginnis/; s3://mcginnislab-gbm-atlas/, region us-east-1, no-sign-request), distributed under CC-BY-4.0. The deposit contains the self-contained interactive HTML viewer, the full per-peak master score table, per-cell-type top-1000 candidate tables, the pseudobulk matrix, the union peak reference, peak-provenance tables, cohort attribution, the schema and changelog, and a regeneration receipt verifying that the shipped snapshot was rebuilt end-to-end before upload. All pipeline source code is at the GitHub repository https://github.com/jpmcginnis/HumanAAVProject/tree/main/glioblastoma-snATAC-seq-pipeline

#### Interactive atlas and candidate downloads

The browsable HTML viewer is mirrored at data.mcginnislab.org.

### AI authorship statement

All code for dataset acquisition, interrogation, harmonization, peak calling, cell-type annotation transfer and integration, cross-cohort reconciliation, replicability and specificity scoring, and figure generation was generated by Anthropic’s Claude Code (Opus 4.7) across numerous interactive sessions, under continual and iterative direction by the authors. The authors specified all scientific choices, including dataset inclusion and exclusion criteria, cohort definitions, units of analysis, cell-type ontology resolution, thresholds for peak replicability and specificity, and biological framing of the results, often in discussion with Claude. The authors reviewed outputs at each step and redirected Claude’s implementation whenever a coding choice conflicted with the intended biological question. The authors take full responsibility for all content.

---

**Supplementary Figure 1.**
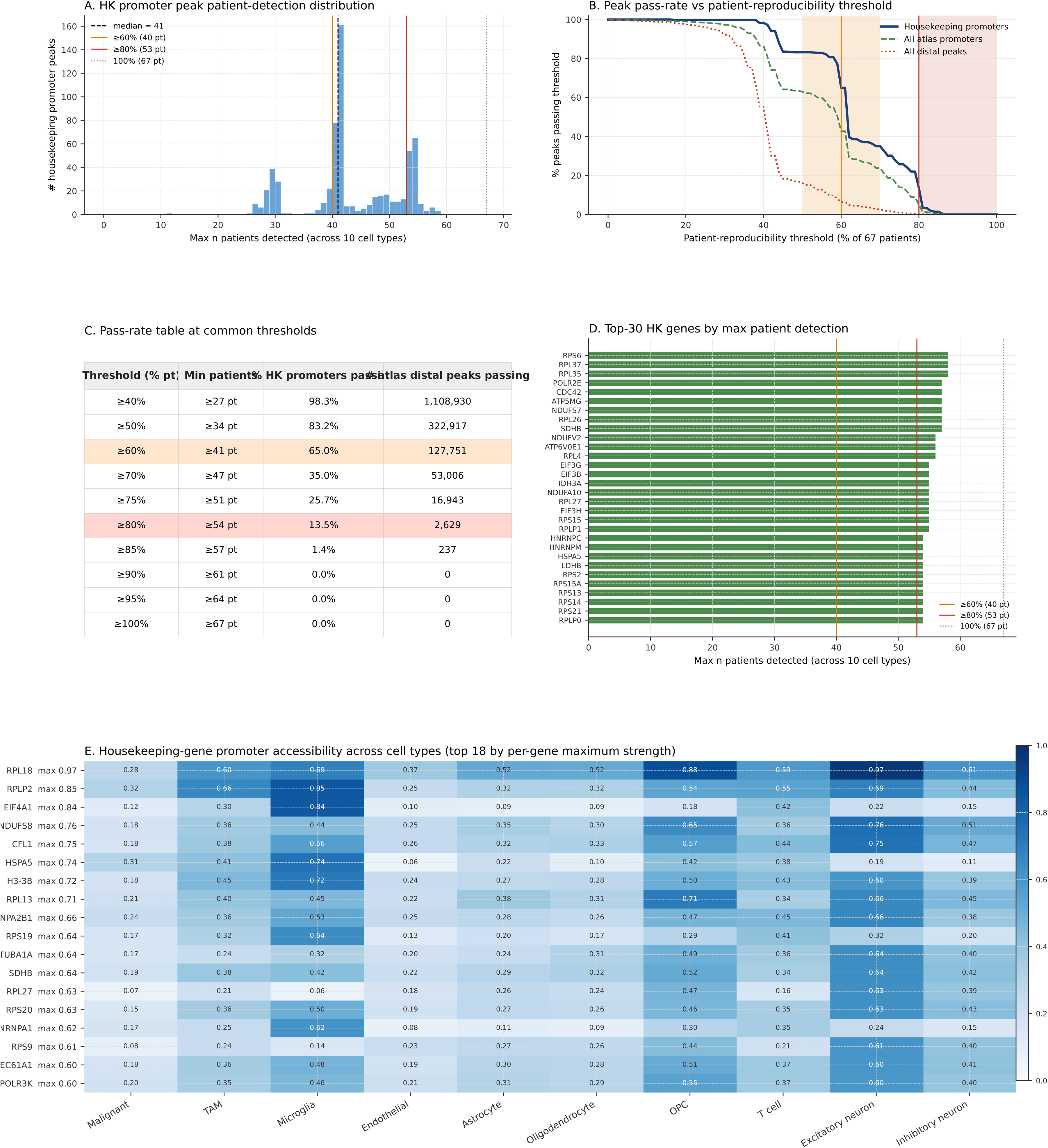
Housekeeping-gene promoter patient-detection sets the technical reproducibility ceiling and motivates the ≥60% patient-reproducibility threshold used throughout the atlas. All panels are computed on the 67 atlas patients with at least one evaluable pseudobulk (≥50 cells per cell type). A peak is counted as detected in a patient if the patient’s best-detecting cell type has at least 1% of its cells carrying a Tn5 fragment at that peak. (A) Histogram of the patient-detection count for the 637 promoter peaks of 206 canonical housekeeping genes (ribosomal subunits, Pol II subunits, translation factors, splicing/hnRNP, cytoskeletal, chaperones, respiratory-chain, proteasome, and classical constitutive markers). Vertical references mark the median, the 60% and 80% patient-detection thresholds, and the 100% ceiling. (B) Pass-rate curve across the 0–100% patient-reproducibility range, for the 637 housekeeping promoters (navy), all atlas promoters ≤2 kb from any TSS (dashed green), and all atlas distal peaks ≥2 kb from any TSS (dotted red). (C) Pass-rate table at ten common reproducibility thresholds, giving the minimum patient count, housekeeping-promoter pass rate, and number of atlas distal peaks meeting each threshold. ≥60% and ≥80% rows highlighted. (D) Top thirty housekeeping genes ranked by maximum patient-detection count across all of each gene’s promoter peaks. (E) Heatmap of housekeeping-gene promoter accessibility across the 10 primary cell types for the top-18 HK genes by per-gene maximum strength.

**Supplementary Figure 2.**
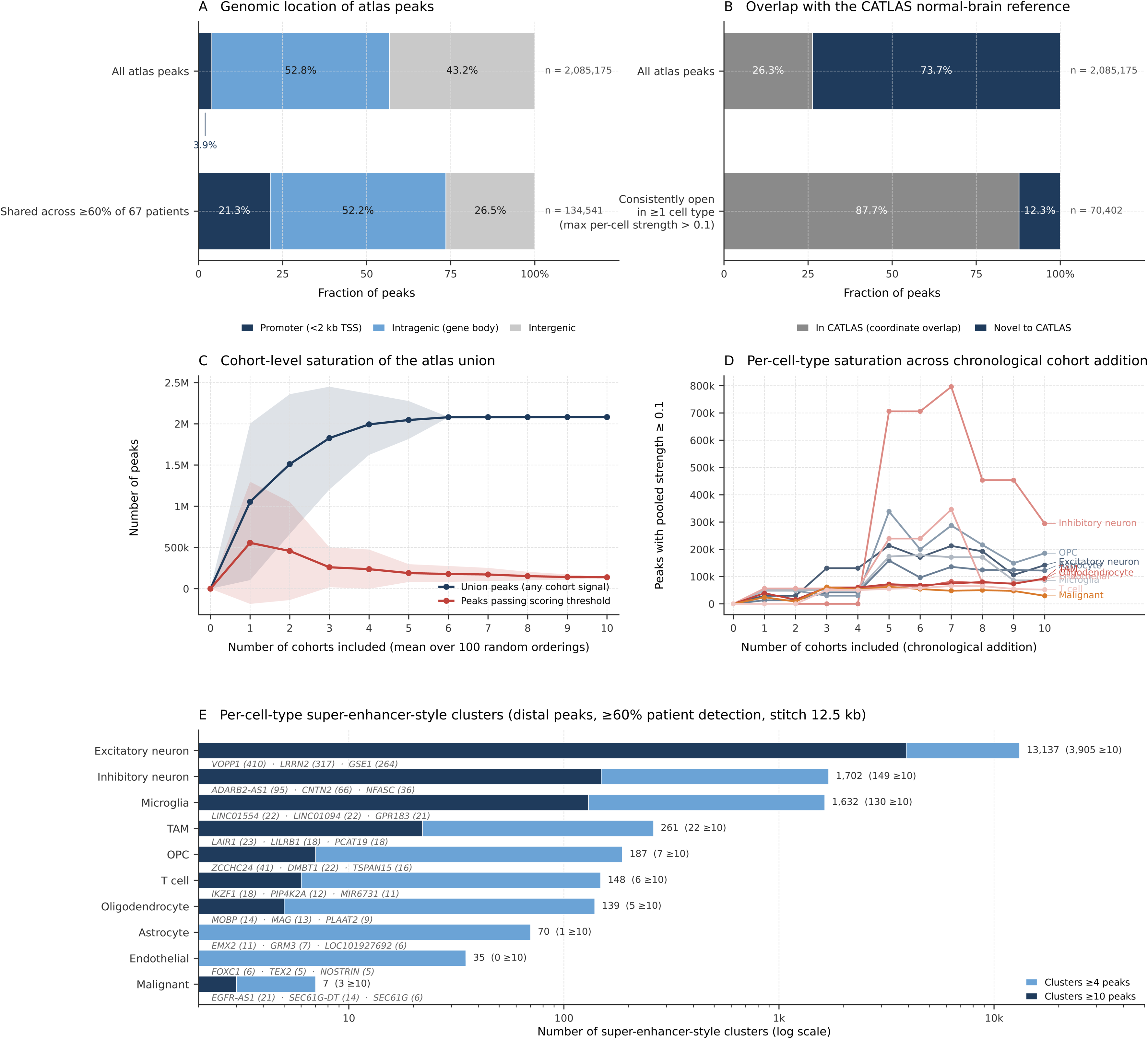
Atlas peak composition, CATLAS overlap, saturation behavior, and super-enhancer-style clustering. (A) Genomic-location breakdown of all 2,085,175 atlas peaks compared to the 134,541 peaks detected in at least 60% of the 67 atlas patients. Promoter-proximal peaks are enriched ∼5-fold in the shared subset (21.3% vs 3.9%). (B) Coordinate overlap between atlas peaks and the CATLAS normal-brain snATAC-seq reference (544,735 peaks). Restricting to peaks that are consistently open in at least one cell type (max per-cell strength > 0.1; 70,402 peaks) raises CATLAS coverage from 26.3% to 87.7%. (C) Rarefaction curve of atlas peak counts as cohorts are randomly added (mean ± SD over 100 random orderings), for both the union of peaks with any cohort signal and the subset passing the atlas scoring threshold. (D) Per-cell-type peak inventory as cohorts are added in chronological order of publication, counting peaks with pooled per-cell-type strength ≥ 0.1 fragments per cell. (E) Per-cell-type super-enhancer-style clusters, defined by stitching distal peaks within 12.5 kb whose per-primary-cell-type patient detection rate reached ≥60%, retaining clusters of ≥4 peaks. Pale bars give the total cluster count per cell type; dark overlay gives the subset with ≥10 peaks. The top three anchor genes by cluster size are listed under each row.

